# A subcortical switchboard for exploratory, exploitatory, and disengaged states

**DOI:** 10.1101/2023.12.20.572654

**Authors:** Mehran Ahmadlou, Maryam Yasamin Shirazi, Pan Zhang, Isaac L. M. Rogers, Julia Dziubek, Sonja B. Hofer

## Abstract

To survive in evolving environments with uncertain resources, animals need to dynamically adapt their behavior and exhibit flexibility in choosing appropriate behavioral strategies, for example, to exploit familiar choices, to explore and acquire novel information, or to disengage altogether. Previous studies have mainly investigated how forebrain regions represent choice costs and values as well as optimal decision strategies during explore/exploit trade-offs. However, the neural mechanisms by which the brain implements alternative behavioral strategies such as exploiting, exploring or disengaging from the environment, remains poorly understood. Here we identify a neural hub critical for flexible switching between behavioral strategies, the median raphe nucleus (MRN). Using cell-type specific optogenetic manipulations, calcium fiber photometry and circuit tracing in mice performing diverse instinctive and learnt behavioral tasks, we found that the MRN’s main cell types, GABAergic, glutamatergic (VGluT2-positive), and serotonergic neurons, have complementary functions and drive exploitation, exploration and disengagement, respectively. Suppression of MRN GABAergic neurons, for instance through inhibitory input from lateral hypothalamus which conveys strong positive valence to the MRN, leads to perseverance in current actions and goals, and thus promotes exploitatory behavior. In contrast, activation of MRN VGluT2+ neurons drives exploratory behavior. Activity of serotonergic MRN neurons is necessary for general task engagement. Input from the lateral habenula conveying negative valence suppresses serotonergic MRN neurons, leading to disengagement. These findings establish the MRN as a central behavioral switchboard, uniquely positioned to flexibly control behavioral strategies. These circuits thus may also play an important role in the etiology and possible treatment of major mental pathologies such as depressive or obsessive-compulsive disorders.

## Main text

All animals are adept at switching their modus operandi to adjust to changes in environmental conditions, the availability of resources and internal needs. At any moment in time, animals have to decide between competing strategies that govern their interactions with environmental resources, such as whether to explore, to exploit, or to disengage from the environment. Exploitation, the efficient utilization of known resources, ensures immediate gains and minimizes risk. In contrast, exploration entails the more labor-intensive endeavor of seeking out novel opportunities and gathering knowledge to increase the chance of future gains^1,2^. Alternatively, animals can disengage from active pursuit of goals with the benefit of conserving energy and minimizing exposure to predator risk. Previous studies on the neural basis of explore/exploit decisions have primarily focused on the role of prefrontal cortical areas (PFC) in evaluating the costs and computing the anticipated value of different choices^3–7^, and on the role of dopaminergic signaling in striatum and PFC for updating expected choice value^5,8,9^. However, maintaining the correct balance between exploratory, exploitatory and disengaged states is crucial for survival across the animal kingdom, indicating that neural circuits underpinning these behavioral strategies are evolutionarily conserved^10–14^. Although the identity of these circuits has remained elusive, we speculated that they must encompass subcortical neural pathways that enable animals to maintain or switch between behavioral strategies independently of higher-order cognitive functions and telencephalic circuits. In this study, we identify the MRN in the brainstem as a key switchboard for controlling exploitation, exploration and disengagement across instinctive and acquired behaviors. The MRN provides a neural nexus at the interface of internal state, affective and cognitive information from brain regions such as the hypothalamus, habenula and prefrontal cortex^15–18^. Alongside the dorsal raphe nucleus, it is a main source of the neuromodulator serotonin, which has been broadly implicated in behavioral flexibility, perseverative behavior and obsessive disorders^13,19–23^. However, the MRN does not only contain serotonergic neurons. In fact, only ∼5-10% of MRN neurons are serotonergic^24,25^, while the majority is GABAergic (VGAT+, ∼60%)^25^ or glutamatergic (VGluT2+, ∼25%^25^ and VGluT3+, mainly overlapping with SERT+^26^).

### VGAT+ MRN neurons regulate perseverance and exploitatory behavior

To test if any of the MRN cell types play a role in regulating animals’ natural behavioral strategies for interacting with the environment, we aimed to establish a paradigm in which mice display exploratory, exploitatory and disengaged behavioral interaction states during instinctive, naturalistic behavior without the need for prior knowledge or training. Freely-moving mice were exposed to twenty small, novel objects, and their behavior was video-captured and scored (**Fig. 1a; movie S1**). Labeled actions during this multi-novel object interaction (MNOI) test included actions attributed to a specific object such as approaching and leaving an object, deep interaction (defined as the mouse grabbing, carrying or biting the object), and sniffing interaction (defined as the mouse sniffing and then leaving an object without deep interaction), as well as sitting and walking without object interaction. We trained an unsupervised hidden Markov model (HMM)^27^ on control mice to categorize the sequences of labeled actions into three interaction states (see Methods). In periods assigned to state 1 by the HMM mice were mainly engaged in deep, long-duration interactions with one or few objects. We named this state in which mice showed such sustained actions exploitatory state. In state 2 mice exhibited rapid switching between several objects without deep interactions, this state was therefore named exploratory state. In state 3, the disengaged state, mice were passive or walked around without object interaction (**Fig. 1b; Supplementary Fig. 1; movie S1**). Control mice spent roughly equal amounts of time in each of these three states (**Fig. 1b**).

**Fig. 1:**
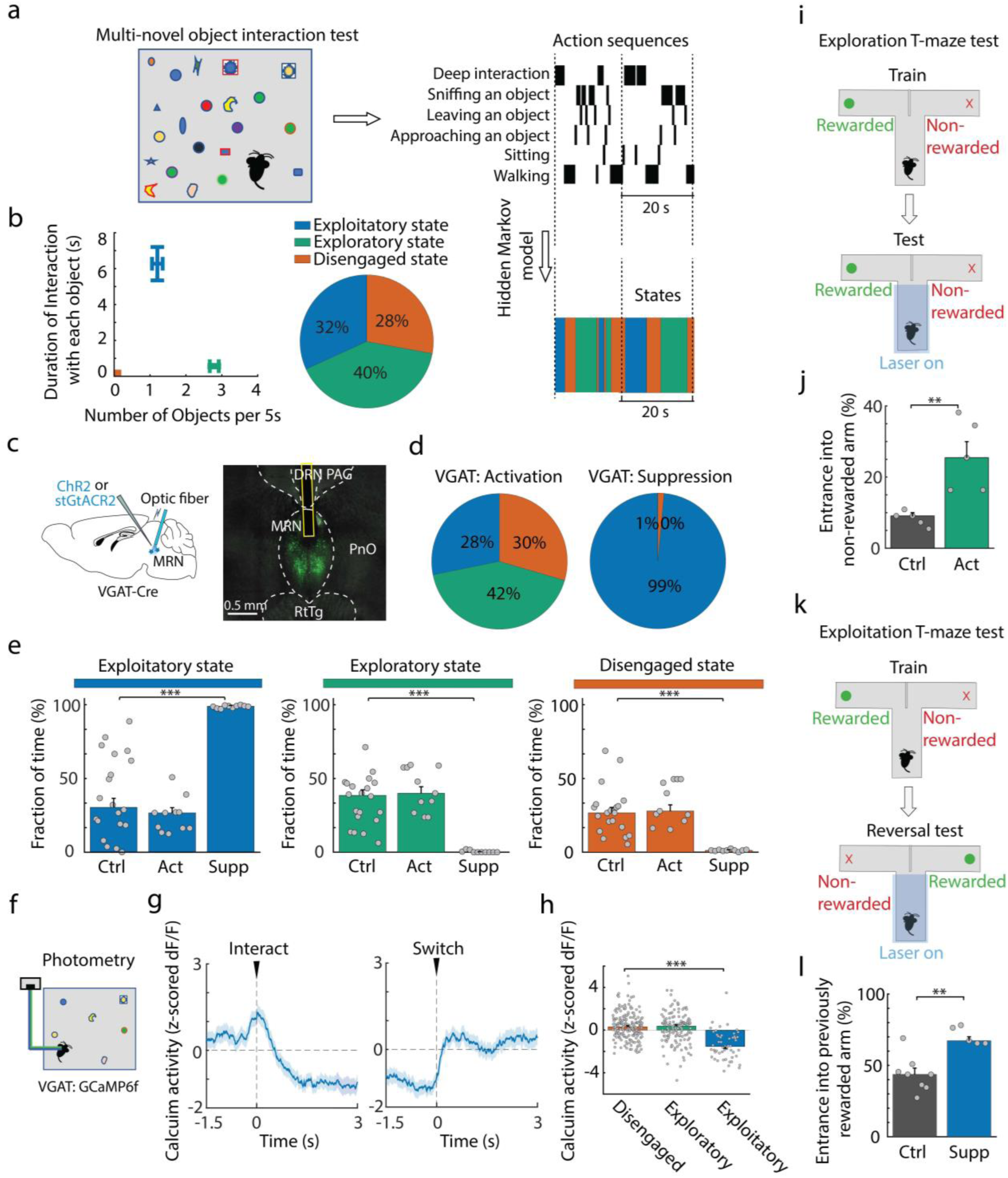
VGAT expressing MRN neurons regulate exploitatory state. **a,** Schematic of the Multi-Novel Object Interaction (MNOI) test (left), and an example sequence of annotated actions (top right), with three interaction states extracted by a Hidden Markov Model (HMM, bottom right). **b,** Left: average duration of interaction with each object plotted against the frequency of interactions with different objects in each HMM state in control mice (median ± bootstrap standard error, N = 20 experiments from 10 mice). Right: Median fraction of time spent in each state. **c,** Left: schematic of optogenetically activating or suppressing VGAT+ MRN neurons, using ChR2 or stGtACR2. Right: example image of virus expression in VGAT+ MRN neurons with optic fiber position. DRN: dorsal raphe nucleus, MRN: median raphe nucleus, PAG: periaqueductal gray, PnO: pontine reticular formation, RtTg: reticulotegmental nucleus. **d,** Median fraction of time spent in the exploitatory (blue), exploratory (green), and disengaged state (orange) during the MNOI test in mice with activation (left) or suppression (right) of VGAT+ MRN neurons. **e,** Fraction of time spent in each interaction state in tdTomato control mice, and mice shown in **d** with activation (act) or suppression (supp) of VGAT+ MRN neurons. Bars indicate median values, error bars are bootstrapped standard error and circles individual experimental sessions. Exploitatory state: ctrl vs act: P = 0.2895, ctrl vs supp: P = 3.1×10^-7^; exploratory state: ctrl vs act: P = 0.4587, ctrl vs supp: P = 1.1×10^-6^; disengaged state: ctrl vs act: P = 0.6774, ctrl vs supp: P = 5.6×10^-5^, two-sided t-test with Bonferroni multi-comparison correction. N = 20, 11 and 10 experiments from 10, 6 and 5 mice in tdTomato, VGAT+ neuron activation and VGAT+ suppression groups. **f,** Schematic of calcium photometry recording from VGAT+ MRN neurons in mice exposed to multiple novel objects. **g,** Average z-scored calcium activity trace (mean ± s.e.m.) of VGAT+ MRN neurons during object interactions aligned either to the onset of deep interactions (left) or the time of switching between objects (right). **h,** Median z-scored calcium activity of the VGAT+ MRN neurons during disengaged states (N = 190 events from 5 mice), exploratory states (N = 139 events, 5 mice, disengaged vs exploratory P = 0.2194) and exploitatory states (N = 51 events, 5 mice, disengaged vs exploitatory: P = 1.5×10^-12^), nested ANOVA with Bonferroni multi-comparison correction. **i,** Schematic of experimental design to quantify levels of exploration during a T-maze test. During the test, in each trial photo-stimulation is applied throughout the central arm until mice turn into the left or right arm. **j,** Percentage of trials with entrance into the non-rewarded arm during the exploration T-maze test, in tdTomato control mice (N = 5 mice) and mice with optogenetic activation of VGAT+ (right, 5 mice, P = 0.0048, two-sided t-test) MRN neurons. **k,** Schematic of experimental design to quantify levels of exploitation during a reversal test in the T-maze. **l,** Percentage of trials with entrance into the previously rewarded arm during the reversal T-maze test, in tdTomato control mice (N = 8 mice) compared with mice with optogenetic suppression of VGAT+ (right; 5 mice, P = 0.0011, two-sided t-test) MRN neurons. *: p-value < 0.05, **: p-value < 0.01, ***: p-value < 0.001.

Because they comprise the largest fraction of neurons in the MRN^25^, we first investigated the potential role of VGAT-positive MRN (MRN^VGAT^) neurons in regulating animals’ interaction states. To test whether manipulating the activity of MRN^VGAT^ neurons changes the time animals spend in an exploratory, exploitatory or disengaged state, we expressed either a Cre-dependent soma-targeted inhibitory opsin, stGtACR2^28^, or an excitatory opsin, ChR2, in MRN^VGAT^ neurons of VGAT-Cre mice via carefully targeted viral injections (**Fig. 1c; Supplementary Fig. 2a**), to optogenetically suppress or activate these neurons, respectively, during the MNOI test. Suppression of MRN^VGAT^ neurons induced a dramatic increase in sustained interactions with individual objects, with much fewer switches between objects compared to control mice expressing tdTomato in MRN neurons (**Fig. 1d,e, movie S2, Supplementary Fig. 2b**). Mice thus spent most time in what we defined as exploitatory state when MRN^VGAT^ neurons were suppressed, and, consequently, the duration of both exploratory and disengaged states was decreased to near zero (**Fig. 1d,e**). Activation of MRN^VGAT^ neurons in the MNOI test did not result in a significant change in the duration of the three interaction states (**Fig. 1d,e**). It did, however, significantly decrease for how long mice deeply interacted with individual objects and how often they switched between objects, suggesting that activation of MRN^VGAT^ neurons may suppress sustained interactions (**Supplementary Fig. 2b,c**). To determine whether MRN^VGAT^ neurons are naturally suppressed during sustained object interactions and exploitatory behavior we recorded population calcium signals of MRN^VGAT^ neurons (after Cre-dependent GCaMP6f expression in VGAT-Cre mice) using fiber photometry during the MNOI test (**Fig. 1f**). In line with the effects of optogenetic manipulation, the activity of MRN^VGAT^ neurons was significantly suppressed throughout sustained object interactions, and returned to baseline levels when animals switched to a different object (**Fig. 1g**). MRN^VGAT^ neuron activity was also decreased in general during the exploitatory state extracted from the HMM model, compared to exploratory and disengaged states (**Fig. 1h**).

The above results show that activity of MRN^VGAT^ neurons can strongly influence how much mice persevere in an interaction. However, it remains unclear how these effects translate to more conventional definitions of exploitatory and exploratory behavior, such as persisting in the pursuit and exploitation of a well-known option for acquiring resources versus the exploration of alternative options. To test if MRN neurons can also regulate explore/exploit choices in tasks in which mice act on previously gained knowledge of how to maximize short-term benefits, we trained food-restricted mice on a T-maze task^29,30^, in which a food reward was provided consistently only in one specific arm and not the other. In well-trained mice (>90% correct trials in two sequential sessions), the percentage of trials in which they enter the non-rewarded arm indicates their tendency to explore (**Fig. 1i**). Interestingly, when MRN^VGAT^ neurons expressing ChR2 were optogenetically activated in well-trained mice in the central arm of the T-maze, mice were more likely to choose the non-rewarded arm, i.e. the exploratory option (**Fig. 1j**). To test if suppression of MRN^VGAT^ neurons instead biases mice towards exploitatory behavior, we reversed the reward location in the T-maze in well-trained mice, i.e. the previously rewarded arm was not rewarded any more (**Fig. 1k**). In this scenario, control mice start exploring the other, previously non-rewarded arm and eventually learn to expect reward only at this location. When stGtACR2-expressing MRN^VGAT^ neurons were optogenetically suppressed in this reversal-learning paradigm in the central arm of the T-maze, mice remained more likely to choose the previously rewarded arm, showing an increase in exploitatory behavior (**Fig. 1l**). This result is highly consistent with the increased duration of exploitatory states caused by MRN^VGAT^ neuron suppression during the MNOI test (**Fig. 1d,e**), demonstrating that during both instinctive and learned goal-directed behaviors, suppression of MRN^VGAT^ neurons causes perseverance towards a current or familiar choice, consistent with an exploitatory state.

### VGLUT2+ MRN neurons drive exploratory behavior

Next, we tested if MRN^VGluT2^ neurons also play a role in regulating animals’ interaction states in the MNOI test. We expressed ChR2 or stGtACR2 in MRN of VGluT2-Cre mice to optogenetically activate or deactivate MRN^VGluT2^ neurons, respectively (**Fig. 2a**). Activating MRN^VGluT2^ neurons in the MNOI test significantly increased the time mice spent in an exploratory state, and specifically decreased the duration of exploitatory states, compared to control mice (**Fig. 2b,c; movie S3**). Additionally, mice switched more often between different objects when MRN^VGluT2^ neurons were activated (**Supplementary Fig. 3**), further indicating increased exploratory behavior. Suppression of MRN^VGluT2^ neurons in the MNOI test did not evoke significant changes in the duration of any of the states (**Fig. 2b,c**), or in how often mice switched between objects (**Supplementary Fig. 3**). In line with the effects of optogenetic manipulation, calcium signals recorded with fiber photometry (**Fig. 2d**) showed that activity of MRN^VGluT2^ neurons significantly increased during the exploratory state and when mice switched from one object to another, but was not different from baseline during deep interactions (**Fig. 2e,f**).

**Fig. 2:**
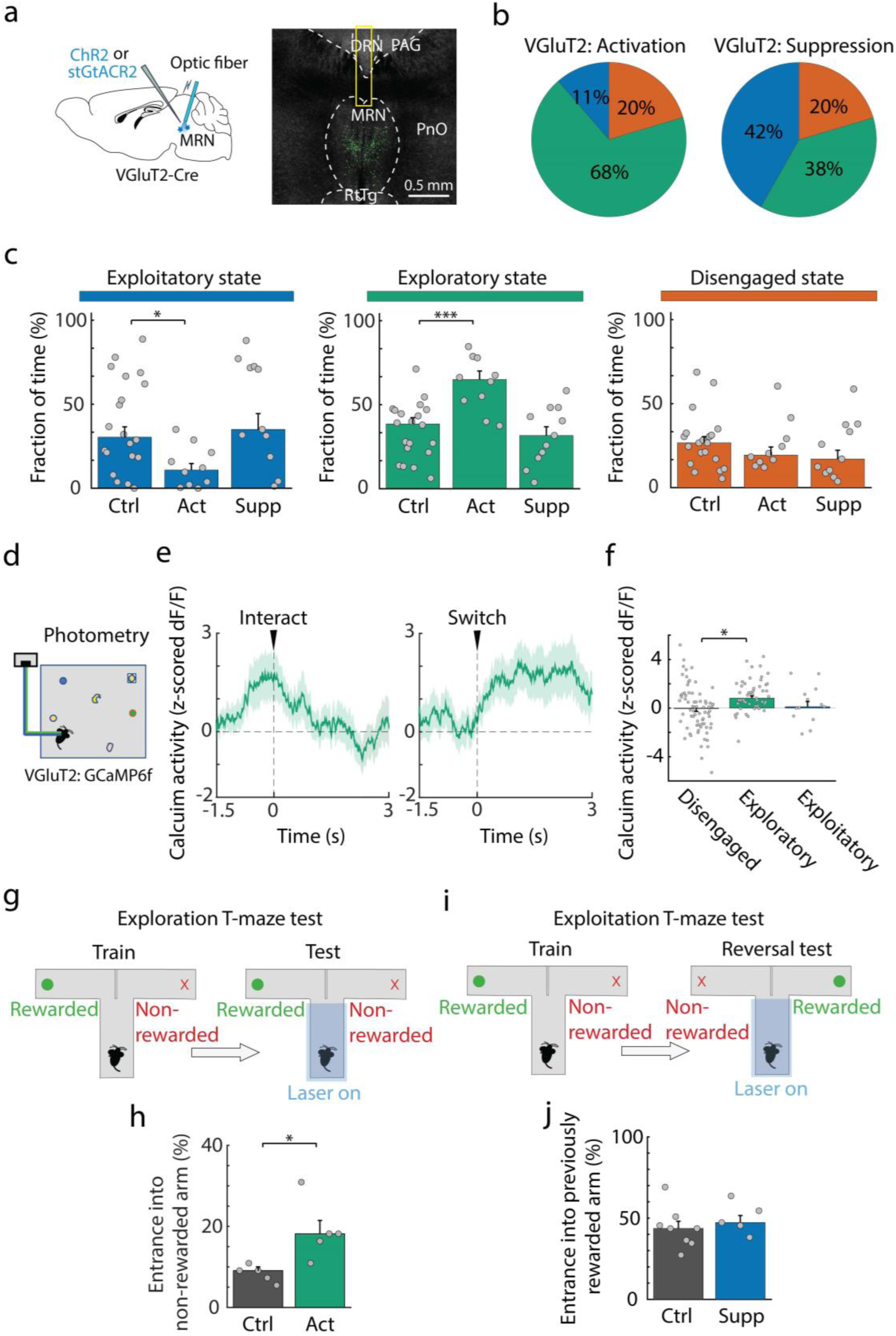
VGluT2 expressing MRN neurons drive exploratory behavior. **a,** Left: schematic of optogenetically activating or suppressing VGluT2+ MRN neurons, using ChR2 or stGtACR2. Right: example image of virus expression in VGluT2+ MRN neurons with optic fiber positions. DRN: dorsal raphe nucleus, MRN: median raphe nucleus, PAG: periaqueductal gray, PnO: pontine reticular formation, RtTg: reticulotegmental nucleus. **b,** Median fraction of time spent in the exploitatory (blue), exploratory (green), and disengaged state (orange) during the MNOI test in mice with activation (left) or suppression (right) of VGluT2+ MRN neurons. **c,** Fraction of time spent in each interaction state in tdTomato control mice, and mice shown in **b**. Exploitatory state: ctrl vs act: P = 0.0216, ctrl vs supp: P = 0.9615; exploratory state: ctrl vs act: P = 0.0004, ctrl vs supp: P > 0.9999; disengaged state: ctrl vs act: P > 0.9999, ctrl vs supp: P = 0.9085, two-sided t-test with Bonferroni multi-comparison correction. N = 20, 10 and 11 experiments from 10, 5 and 5 mice in tdTomato, VGluT2+ excitation and VGluT2+ inhibition groups. **d,** Schematic of calcium photometry recording from VgluT2+ MRN neurons in mice exposed to multiple novel objects. **e,** Average z-scored calcium activity trace (mean ± s.e.m.) of VGluT2+ MRN neurons during object interactions aligned either to the onset of deep interactions (left) or the time of switching between objects (right). **f,** Median z-scored calcium activity of the VGluT2+ MRN neurons during disengaged, exploratory and exploitatory states. N = 72 disengaged, 53 exploratory and 13 exploitatory state events from 3 mice. Disengaged vs exploratory: P = 0.0122, disengaged vs exploitatory: P = 0.9272, nested ANOVA with Bonferroni multi-comparison correction. **g,** Schematic of experimental design to quantify levels of exploration during a T-maze test. During the test, in each trial photo-stimulation is applied throughout the central arm until mice turn into the left or right arm. **h,** Percentage of trials with entrance into the non-rewarded arm during the T-maze test, in tdTomato control mice (N = 5 mice) and mice with optogenetic activation of VGluT2+ (left, 5 mice, P = 0.0149, two-sided t-test) MRN neurons. **i,** Schematic of experimental design to quantify levels of exploitation during a reversal test in the T-maze. **j,** Percentage of trials with entrance into the previously rewarded arm during the reversal T-maze test, in tdTomato control mice (N = 8 mice) compared with mice with optogenetic suppression of VGluT2+ (left; 5 mice, P = 0.3873, two-sided t-test) MRN neurons. *: p-value < 0.05, ***: p-value < 0.001. In Panels c, h and j, bars indicate median values, error bars are bootstrapped standard error and circles individual experimental sessions.

We again used the T-maze task to test if MRN^VGluT2^ neuron activation also promotes exploratory choices in a learned task in which food-deprived mice act on previously gained knowledge. After mice learned that food reward was provided consistently only in one specific arm and not the other, MRN^VGluT2^ neurons expressing ChR2 were optogenetically activated in well-trained mice in the central arm of the T-maze (**Fig. 2g**). During this manipulation, mice were more likely to choose the non-rewarded arm, i.e. the exploratory option (**Fig. 2h**), consistent with the increase in exploratory behavior when MRN^VGluT2^ neurons were activated in the MNOI test (**Fig. 1b,c**). However, suppression of MRN^VGluT2^ neurons did not in turn induce a bias towards exploitatory choices, as mice did not choose the previously rewarded arm more often than control animals when the reward location was reversed (**Fig. 2i,j**). This was again consistent with the lack of effect of MRN^VGluT2^ neuron suppression on behavior during the MNOI test (**Fig. 2b,c**).

These findings indicate that activation of MRN^VGluT2^ neurons promotes exploratory choices during both instinctive and learned goal-directed behaviors. Intriguingly, activation of MRN^VGAT^ neurons had also increased exploratory choices in the T-maze test (**Fig. 1i,j**). However, this test with two forced choices cannot distinguish if these manipulations actively drive exploratory choices or merely prevent perseverance and thus reduce exploitatory behavior. To discern these possibilities, we assessed the effect of activating either MRN^VGluT2^ or MRN^VGAT^ neurons during another behavioral task, a nose-poke reward association task with distractors (**Supplementary Fig. 4a**). In this task, water-restricted mice learned to first insert their snout into a nose port to then receive a water reward upon licking the water port on the opposite wall. The other walls held additional nose ports which were irrelevant for the task. Optogenetic activation of MRN^VGluT2^ neurons biased mice away from the task, towards exploration of the task-irrelevant nose ports (**Supplementary Fig. 4b**). Activation of MRN^VGAT^ neurons, in contrast, had no effect on behavior and did not evoke interactions with the task-irrelevant nose ports (**Supplementary Fig. 4c**), indicating that only activity of MRN^VGluT2^ neurons actively drives exploratory behavior.

### VGAT+ and VGLUT2+ MRN neurons modulate affective state

Changes in arousal level and valence – the affective signed value associated with a stimulus or context, are crucial for the manifestation of motivational states^31^, and therefore likely play an important role in regulating interactions with the environment. Next, we thus aimed to determine to what degree changes in affective state can explain the effects of MRN^VGluT2^ and MRN^VGAT^ neuron manipulation on explore/exploit decisions. We used a self-stimulation test, commonly used to assess the valence of a neural manipulation^32,33^, in which mice are presented with two nose-poke ports, only one of which triggers optogenetic self-stimulation upon entering (opto-linked port, **Fig. 3a**). Suppression of MRN^VGAT^ neurons led to a very strong preference for the opto-linked port, indicating that this manipulation is reinforcing and induces strong positive valence (**Fig. 3b**). The positive valence conveyed by MRN^VGAT^ neuron suppression was so dominant that it could even override strongly aversive cues. When normal mice were presented with an object coated with an innately highly aversive substance, Trimethylthiazoline (TMT; a constituent of fox urine)^34^, they usually carefully approached the TMT object only a few times, often followed by a fast retreat (‘escape’, see Methods, **Fig. 3c-e, movie S4**). Suppression of MRN^VGAT^ neurons dramatically increased how often mice approached the TMT object, prevented escapes from it (**Fig. 3d,e**), and, remarkably, even elicited deep interactions with the normally highly aversive object (**movie S5**), an action never observed in normal mice. These findings show that suppression of MRN^VGAT^ neurons induces perseverance in the current choice or goal by bestowing highly positive valence, overriding other motives.

**Fig. 3:**
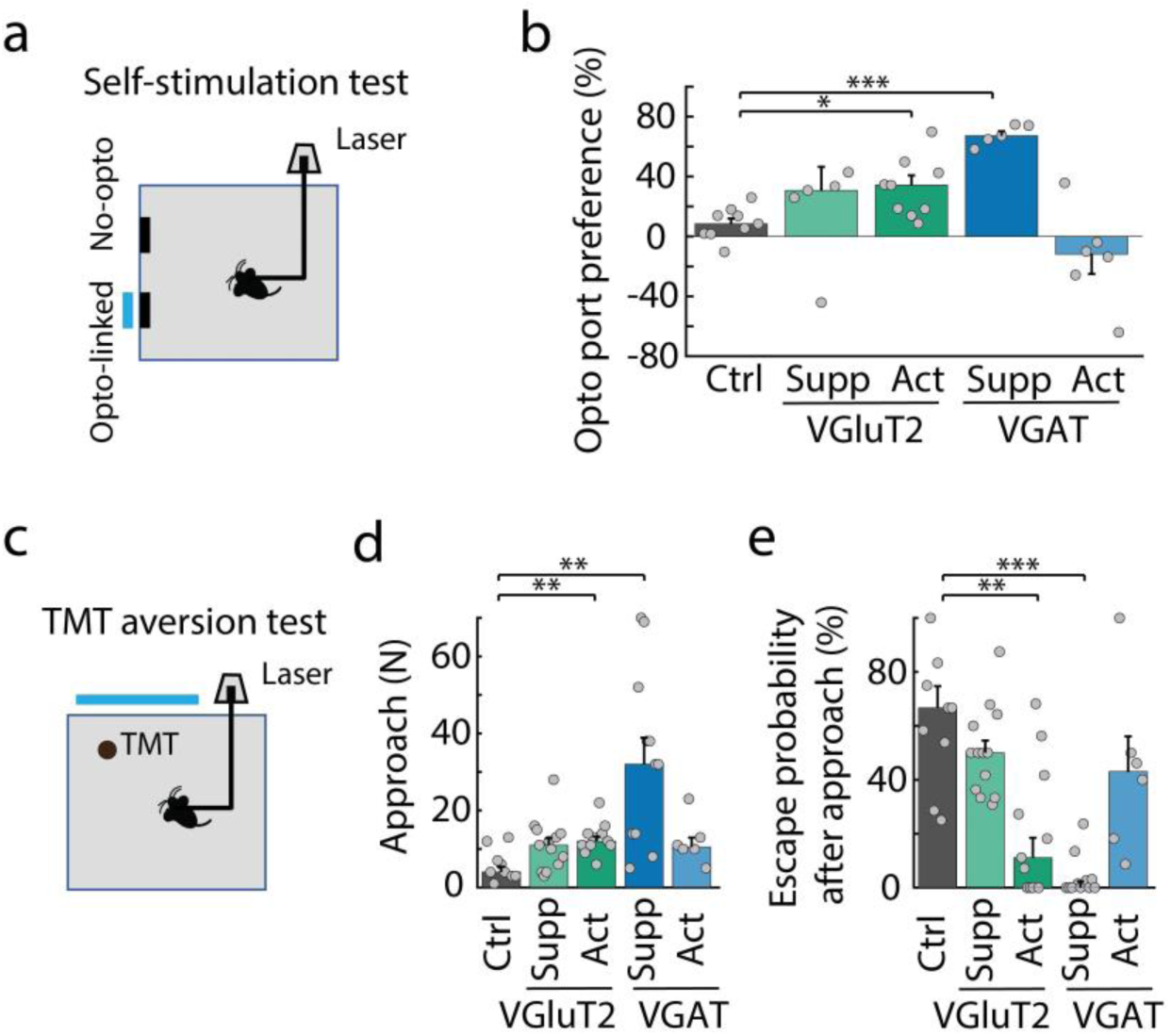
Effect of manipulation of VGAT and VGluT2 expressing MRN neurons on valence. **a,** Schematic of the experimental design for a self-stimulation test. **b,** Preference for entering the opto-linked nose-poke port (100 × (number of times entries into the opto-linked nose-poke port - number of times entries into the non-stimulation nose-poke port)/total number of entries into both nose-poke ports) in tdTomato mice and mice with activation or suppression of VGAT+ and VgluT2+ MRN neurons. Control mice (N = 9) compared to mice with VGluT2+ neuron suppression (N = 5 mice, P > 0.9999), VGluT2+ neuron activation (N = 9 mice, P = 0.0248), VGAT+ suppression (N = 5 mice, P = 4.5×10^-7^) and VGAT+ neuron activation (N = 6 mice, P = 0.2912); two-sided t-test with Bonferroni multi-comparison correction. **c,** Schematic of the TMT aversion test. **d,** Number of approaches of the TMT- coated object. Control mice compared to mice with VGluT2+ suppression (P = 0.2126), VGluT2+ activation (P = 0.0074), VGAT+ suppression (P = 0.0080) and VGAT+ activation (P = 0.1413); two-sided t-test with Bonferroni multi-comparison correction. **e,** Escape probability after approaching a TMT-coated object. Control mice (N = 9) compared to mice with VGluT2+ suppression (N = 13 mice, P > 0.9999), VGluT2+ activation (N =11, P = 0.0058), VGAT+ suppression (N =11, P = 2.2×10^-6^) and VGAT+ activation (N = 6, P > 0.9999); two-sided t-test with Bonferroni multi-comparison correction. *: p-value < 0.05, **: p-value < 0.01, ***: p-value < 0.001. In Panels b, d and e, bars indicate median values, error bars are bootstrapped standard error and circles individual experimental sessions.

We repeated the above tests with either activation of MRN^VGluT2^ neurons or suppression of MRN^VGAT^ neurons that had elicited similar results in the T-maze test (**Figs. 1i,j, 2g,h**). Different to a previous study^25^, we found that activation of MRN^VGluT2^ neurons was not aversive (**Supplementary Fig. 5a-c**), but conveyed positive valence, as it increased mice’s preference for the opto-linked port in the self-stimulation test (**Fig. 3b**), and increased the number of approaches of and decreased escape probability from the aversive TMT object (**Fig. 3d,e**). Interestingly, activation of MRN^VGluT2^ neurons also induced heightened levels of locomotion (**Supplementary Fig. 5d,e**) and increased animals’ arousal as measured from changes in pupil size and whisking activity (**Supplementary Fig. 5f-i**). In contrast, activation of MRN^VGAT^ neurons did not induce positive valence nor increase arousal or locomotion (**Fig. 3b,d,e; Supplementary Fig. 5d-i**), providing further evidence that this manipulation does not induce an active state of exploration.

### SERT+ MRN neuron activity is necessary for task engagement

The MRN is a main source of the neuromodulator serotonin, commonly implicated in behavioral flexibility and perseverative behavior^13,19–23^. We therefore tested next if MRN serotonergic (MRN^SERT^) neurons also play a role in balancing exploratory and exploitatory choices. We expressed inhibitory opsin stGtACR2 or excitatory opsin ChR2 specifically in MRN^SERT^ neurons of SERT-Cre mice^35^ (**Supplementary Fig. 6a**), to suppress or activate these neurons, respectively, during the MNOI test, and to quantify the effect of MRN^SERT^ neuron manipulation on the duration of exploitatory, exploratory and disengaged states (**Fig. 4a**). Suppression of MRN^SERT^ neurons strongly increased the time mice spent in a disengaged state and decreased the duration of exploitatory states, compared to control mice (**Fig. 4b,c**). In contrast, optogenetic activation of these neurons had no effect on how long mice spent in each of the states (**Fig. 4b,c**).

**Fig. 4:**
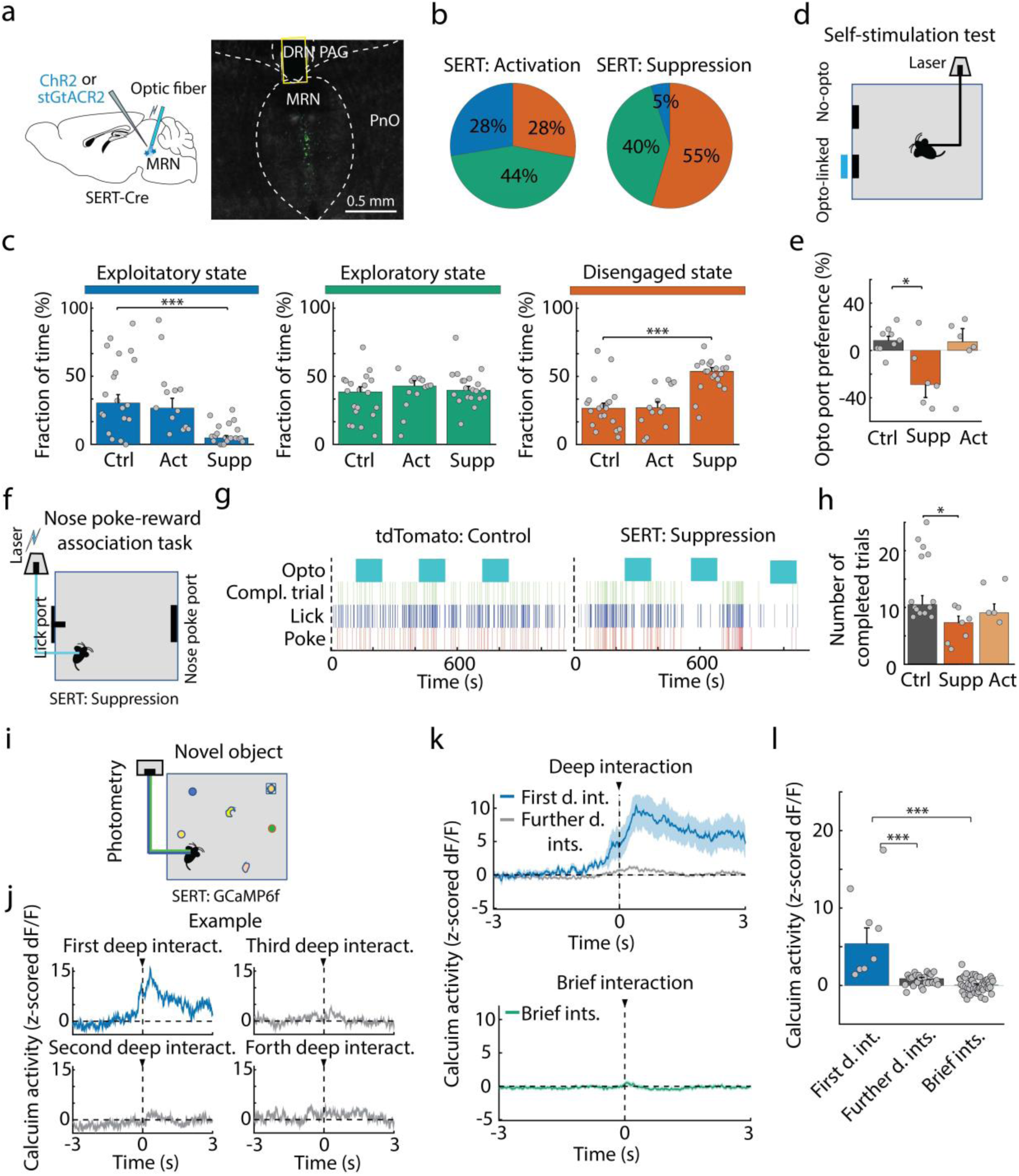
Activity in SERT expressing MRN neurons is necessary for task engagement. **a,** Left: schematic of the experimental design of optogenetically activating or suppressing serotonergic (SERT+) MRN neurons. Right: example image of virus expression in SERT+ MRN neurons with optic fiber position. DRN: dorsal raphe nucleus, MRN: median raphe nucleus, PAG: periaqueductal gray, PnO: pontine reticular formation. **b,** Median fraction of time spent in each interaction state during the MNOI test in mice with activation (left) or suppression (right) of SERT+ MRN neurons. **c,** Fraction of time spent in the exploitatory (blue), exploratory (green), and disengaged state (orange) in tdTomato control mice, and mice shown in **b.** Exploitatory state: control (ctrl) vs activation (act): P > 0.9999; ctrl vs suppression (supp): P = 0.0001; exploratory state: ctrl vs act: P = 0.9897, ctrl vs supp: P = 0.3209; disengaged state: ctrl vs act: P > 0.9999; ctrl vs supp: P = 1.8×10^-5^, two-sided t-test with Bonferroni multi-comparison correction. N = 20, 13 and 20 experiments from 10, 4 and 8 mice in tdTomato, SERT+ activation and SERT+ suppression groups. Bars indicate median and circles depict individual experiments. **d,** Schematic of the experimental design for a self-stimulation test. **e,** Preference for entering the opto-linked nose-poke port (100 × (number of entries into the opto-linked nose-poke port - number of entries into the non-stimulation nose-poke port) / total number of entries into both nose-poke ports) in control mice and mice with activation or suppression of SERT+ MRN neurons. Control mice (N = 9) compared to mice with SERT+ neuron suppression (N = 6 mice, P = 0.0151) and SERT+ neuron activation (N = 6 mice, P > 0.9999); two-sided t-test with Bonferroni multi-comparison correction. Bars depict median and circles indicate individual experiments. **f,** Schematic of experimental design of the nose-poke reward association task. **g,** Examples of task performance (including timing of nose pokes, licks and completed (compl.) trials) of a tdTomato control mouse and a mouse with optogenetic suppression of SERT+ MRN neurons. Blue squares indicate photo-stimulation time periods. **h,** Number of completed trials during photo-stimulation in tdTomato control mice and mice with suppression or activation of SERT+ MRN neurons. N = 14, 7 and 5 mice, respectively; ctrl vs supp: P = 0.0183, ctrl vs act: P = 0.6821; two-sided t-test with Bonferroni multi-comparison correction. Bars indicate median and circles depict individual experiments. **i,** Schematic of calcium fiber photometry recording from SERT+ MRN neurons in mice exposed to multiple objects. **j,** Example traces of z-scored calcium activity of SERT+ MRN neurons, aligned to the onset of object interactions, for the first (blue), second, third and forth (gray) deep interaction with the same object. **k,** Average z-scored calcium activity trace (mean ± s.e.m.) during object interactions for first (top, blue) and following deep interactions (top, gray), and during brief interactions (bottom left, green). N = 4 mice. **l,** Median z-scored calcium response of SERT+ MRN neurons during the first deep interaction (N = 8 events) compared with following deep interactions with the same object (N = 25 events, first vs following deep interactions: P = 5.4×10^-13^, nested ANOVA with Bonferroni multi-comparison correction), and brief interactions (N = 52 events, first deep vs brief interactions: P = 1.0×10^-10^) from 4 mice; two-sided t-test with Bonferroni multi-comparison correction. *: p-value < 0.05, ***: p-value < 0.001. In Panels c, e and h, bars indicate median values, error bars are bootstrapped standard error and circles individual experimental sessions.

In the self-stimulation test, suppression of MRN^SERT^ neurons decreased the animals’ preference for the opto-linked port, compared to control animals (**Fig. 4d,e**), indicating that it conveys negative valence. However, MRN^SERT^ neuron suppression had no effect on measures of arousal (**Supplementary Fig. 6b-f**). Next, we examined whether suppression of MRN^SERT^ neurons also causes disengagement during learned goal-directed behaviors. To this end, we optogenetically suppressed these neurons during a nose-poke reward association task, in which water-restricted mice had learned to first poke into a nose port to subsequently receive a water reward upon licking a water port on the opposite wall (in this case without distractor ports, **Fig. 4f**). In this task, the frequency of completed trials indicates the level of engagement of the animal. When suppressing MRN^SERT^ neurons, animals completed significantly fewer trials compared to control mice (**Fig. 4g,h; movie S6**), confirming that activity of MRN^SERT^ neurons is necessary for task engagement. In contrast, activation of MRN^SERT^ neurons had no effect on engagement levels (**Fig. 4g,h**).

We recorded calcium signals using fiber photometry after expression of Cre-dependent GCaMP6f in MRN of SERT-Cre mice to test how activity of MRN^SERT^ neurons is modulated when encountering different objects (**Fig. 4i**). Activity of MRN^SERT^ neurons was strongly increased during the animals’ first deep interactions with a novel object. Interestingly, further deep or brief interactions with the same item did not cause a calcium response (**Fig. 4j-l**), suggesting that increased MRN^SERT^ neuron activity indicates novelty or salience of an item. We observed a similar pattern of activation when mice interacted with food items instead of novel objects (**Supplementary Fig. 6g,i**). However, interactions with novel objects of negative valence – objects coated in aversive TMT – did not evoke changes in MRN^SERT^ neuron activity (**Supplementary Fig. 6g,h**). This suggests that, different to serotonergic neurons in the dorsal raphe nucleus which have been shown to be activated by both positive and negative valence^36^, MRN^SERT^ neurons respond to salience with positive, but not negative, valence. Our results show that MRN^SERT^ salience signals are necessary for task engagement and sustained interaction with resources. While MRN^SERT^ neuron activation on its own is not sufficient to induce task engagement or sustained interactions, suppression of MRN^SERT^ neurons causes disengagement, potentially by decreasing salience and endowing current choices with negative valence.

Altogether, these findings indicate that MRN^VGAT^ neuron suppression and MRN^VGluT2^ neuron activation drive exploitatory and exploratory choices, respectively. In contrast, MRN^SERT^ activity does not regulate explore/exploit decisions but is necessary for engagement with environmental resources and the continuing pursuit of goals.

### LHb conveys negative valence signals to the MRN, promoting disengagement

Next, we aimed to identify brain areas upstream of MRN that convey information relevant for the generation of exploitatory, exploratory or disengaged states. Using retrograde virus tracing, we identified the lateral hypothalamus (LHA) and lateral habenula (LHb) as two major inputs to the MRN (**Figures 5a,6a**). Because LHb is thought to be associated with negative affective state and depressive-like behaviors^37,38^, we hypothesized that LHb may convey negative valence signals to MRN, important for regulating engagement levels.

**Fig. 5:**
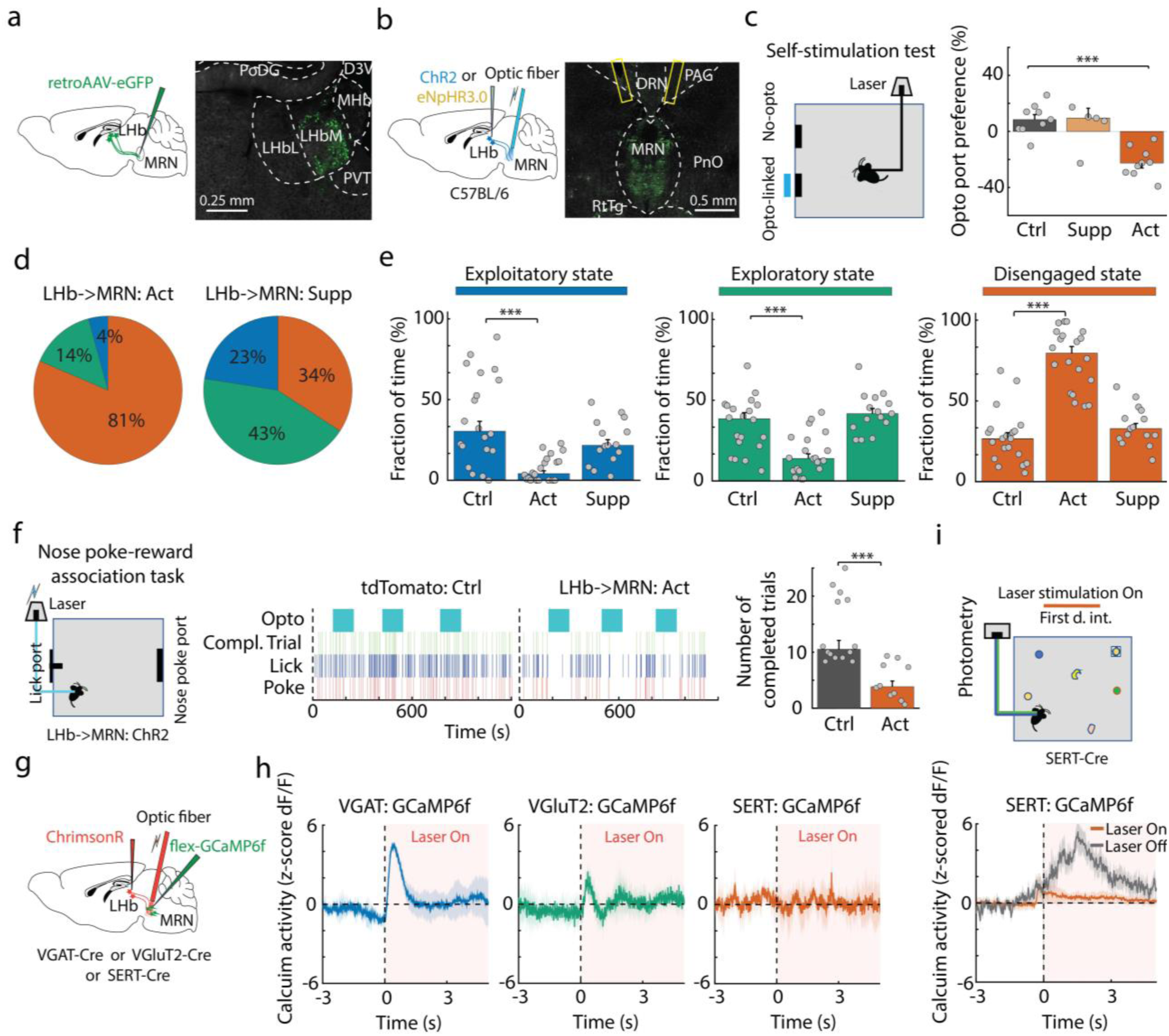
LHb input to MRN modulates the disengaged state. **a,** Schematic of retrograde tracing from MRN using retroAAV (left), and example image of retrogradely labeled neurons in lateral habenula (LHb, right). D3V: dorsal 3rd ventricle, LHbL: lateral part of lateral habenula, LHbM: medial part of lateral habenula, MHb: medial habenula, PoDG: polymorph layer of the dentate gyrus, PVT: paraventricular nucleus of thalamus. **b,** Schematic of optogenetic activation or suppression of LHb input to MRN, using ChR2 or eNpHR3.0 (left), and an example image of ChR2-expressing LHb axons in the MRN with optic fiber positions (right). DRN: dorsal raphe nucleus, MRN: median raphe nucleus, PAG: periaqueductal gray, PnO: pontine reticular formation, RtTg: reticulotegmental nucleus. **c,** Schematic of the self-stimulation test (left), and preference for the opto-linked nose-poke port (100 × (number of entries into the opto-linked nose-poke port - number of entries into the non-stimulation nose-poke port) / total number of entries into both nose-poke ports) in control mice compared to mice with suppression or activation of LHb input to the MRN (right; N = 9, 5 and 9 mice, respectively; P > 0.9999 and P = 5.0×10^-5^ for comparing control mice to activation and suppression of LHb input, respectively; two-sided t-test with Bonferroni multi-comparison correction). Bars depict median and circles indicate individual experiments. **d,** Median fraction of time spent in the three interaction states during the MNOI test in mice with activation (left) and suppression (right) of LHb input to the MRN. **e,** Fraction of time spent in each interaction state in control mice and mice shown in **d** (exploitatory state: ctrl vs act: P = 0.0001, ctrl vs supp: P = 0.1512; exploratory state: ctrl vs act: P = 0.0009, ctrl vs supp: P = 0.3266; disengaged state: ctrl vs act: P = 2.3×10^-10^, ctrl vs supp: P = 0.3697, two-sided t-test with Bonferroni multi-comparison correction). N = 20, 20 and 15 experiments from 10, 9 and 5 mice in control, activation and suppression groups, respectively. **f,** Schematic of nose-poke reward association task with photo-activation of LHb input to the MRN (left) and examples of task performance (including timing of nose pokes, licks and completed (compl.) trials) of a control mouse and a mouse with photo-activation of the LHb input to the MRN. The light blue indicates the photo-stimulation time periods. Bar graph shows the average number of completed trials during photo-stimulation in tdTomato mice and mice with optogenetic activation of LHb input in the MRN (N = 14 and 10 mice, respectively; P = 0.0002, Mann-Whitney U-test). **g,** Schematic of experimental design for recording calcium signals in MRN cell types using fiber photometry, while optogenetically activating LHb input to the MRN in head-fixed mice. **h,** Z-scored calcium traces (mean ± s.e.m.) aligned to laser onset, showing the effect of optogenetic activation of LHb input on the activity of VGAT+ (left), VGluT2+ (middle) and SERT+ (right) neurons in MRN in head-fixed mice. **i,** Schematic of experimental design: calcium recordings from SERT+ MRN neurons in freely moving mice, while activating LHb input to the MRN during the first deep interaction with objects (top). Z-scored calcium activity (mean ± s.e.m.) aligned to onset of first deep object interaction with and without optogenetic activation of LHb input to the MRN (bottom). N = 10 (laser off) vs. 7 (laser on) events from 3 mice; P = 0.0025; two-sided t-test. ***: p-value < 0.001. In Panels c, e and f, bars indicate median values, error bars are bootstrapped standard error and circles individual experimental sessions.

**Fig. 6:**
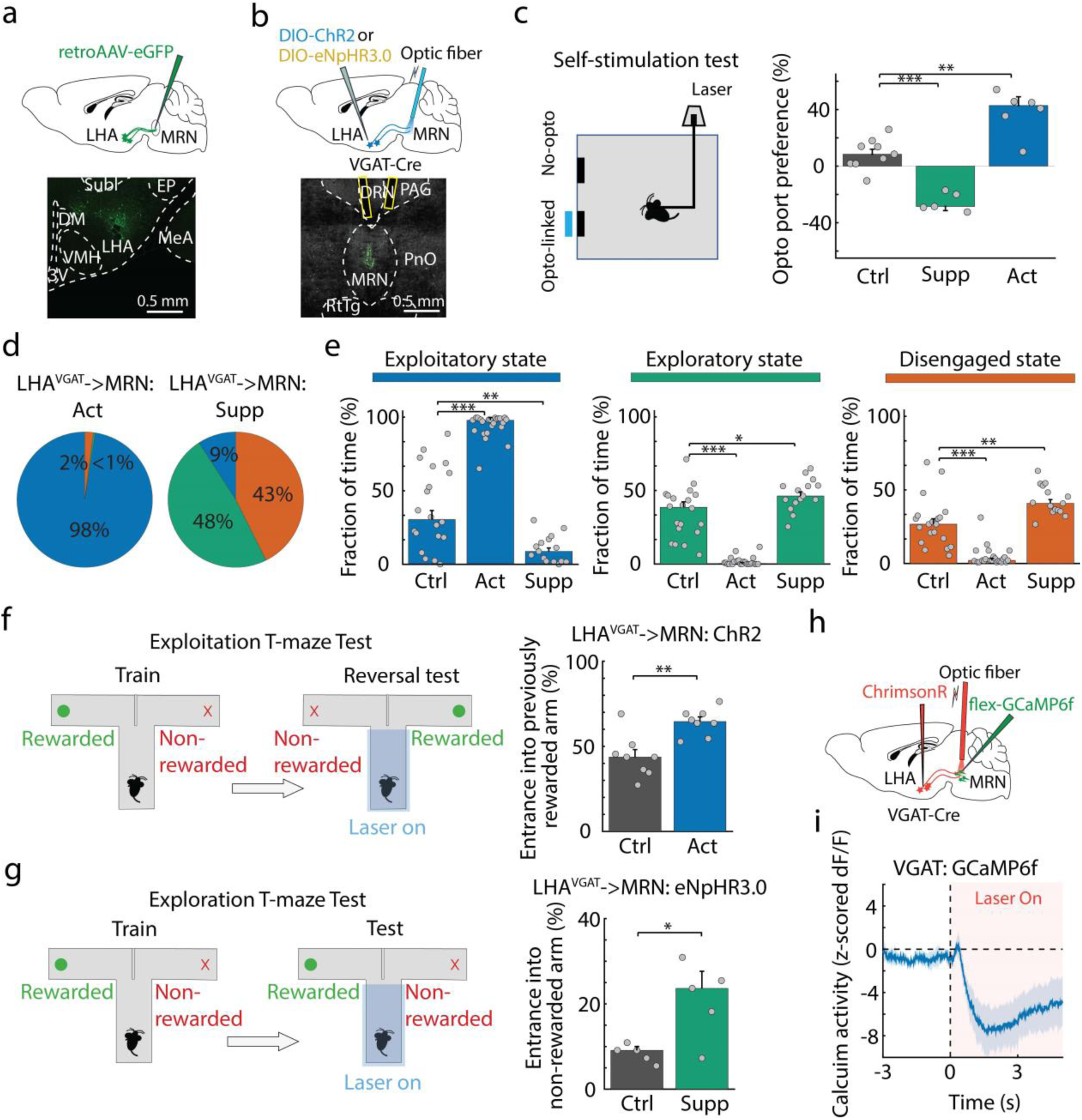
LHA input to MRN bidirectionally regulates exploratory and exploitatory state. **a,** Schematic of retrograde tracing from MRN using retroAAV (top), and example image of retrogradely labeled neurons in lateral hypothalamic area (LHA, bottom). 3V: 3^rd^ ventricle, DM: dorsomedial hypothalamus, EP: entropeduncular nucleus, LHA: lateral hypothalamic area, MeA: medial amygdala, Subl: subincertal nucleus, VMH: ventromedial hypothalamus. **b,** Schematic of optogenetic activation or suppression of LHA VGAT+ input to MRN, using ChR2 or eNpHR3.0 (top), and an example image of LHA axons in MRN with optic fiber positions (bottom). DRN: dorsal raphe nucleus, MRN: median raphe nucleus, PAG: periaqueductal gray, PnO: pontine reticular formation, RtTg: reticulotegmental nucleus. **c,** Schematic of the self-stimulation test (left), and preference for the opto-linked nose-poke port (100 × (number of entries into the opto-linked nose-poke port - number of entries into the non-stimulation nose-poke port) / total number of entries into both nose-poke ports) in control mice compared to mice with suppression or activation of LHA VGAT+ input to the MRN (right; P = 5.0×10^-5^ and P = 0.0012 for comparing control mice to activation and suppression of LHA, respectively; two-sided t-test with Bonferroni multi-comparison correction). **d,** Median fraction of time spent in the exploitatory (blue), exploratory (green), and disengaged state (orange) during the MNOI test in mice with activation (left) and suppression (right) of the LHA VGAT+ input to the MRN. **e,** Fraction of time spent in the three interaction states in control mice, and in mice shown in **d.** Exploitatory state: control (ctrl) vs activation (act): P = 2.1×10^-11^; ctrl vs suppression (supp): P = 0.0015; exploratory state: ctrl vs act: P = 2.3×10^-11^, ctrl vs supp: P = 0.0327; disengaged state: ctrl vs act: P = 4.9×10^-7^; ctrl vs supp: P = 0.0076, two-sided t-test with Bonferroni multi-comparison correction. N = 20, 23 and 15 experiments from 10, 9 and 8 mice in control mice, LHA VGAT+ input activation and LHA VGAT+ input suppression groups. **f,** Left: schematic of experimental design to quantify levels of exploitation during a reversal test in the T-maze. Right: percentage of trials with entrance into the previously rewarded arm during the reversal T-maze test, in control mice (N = 8) compared to mice with optogenetic activation of LHA VGAT+ input to the MRN (8 mice, P = 0.0016, two-sided t-test). **g,** Left: schematic of experimental design to quantify levels of exploration during a T-maze test. During the test, in each trial photo-stimulation is applied throughout the central arm until mice turn into the left or right arm. Right: percentage of trials with entrance into the non-rewarded arm during the exploration T-maze test, in control mice (N = 5) and mice with optogenetic suppression of LHA VGAT+ input to the MRN (5 mice, P = 0.0148, two-sided t-test). **h,** Schematic of experimental design for recording calcium signals in MRN VGAT+ neurons, while optogenetically activating LHA VGAT+ input to the MRN. **i,** Z-scored calcium traces (mean ± s.e.m.) aligned to laser onset, showing the effect of optogenetic activation of LHA VGAT+ input on the activity of the VGAT+ MRN neurons. *: p-value < 0.05, **: p-value < 0.01, ***: p-value < 0.001. In Panels c, e, f and g, bars indicate median values, error bars are bootstrapped standard error and circles individual experimental sessions.

We tested this hypothesis by injecting AAVs for expression of optogenetic constructs in LHb and optogenetically activating either ChR2-expressing LHb axons in MRN, or specifically suppressing this axonal pathway using inhibitory opsin eNpHR3.0 for axonal silencing during the different behavioral paradigms introduced above (**Fig. 5b**). In the self-stimulation test a real-time place preference test, activation of LHb→MRN input significantly decreased mice’s preference for the opto-linked port and the opto-linked chamber (**Fig. 5c; Supplementary Fig. 7a,b**), while silencing the LHb→MRN pathway increased preference for the opto-linked chamber (**Supplementary Fig. 7b**). Furthermore, chronic activation of LHb→MRN input resulted in anhedonia, as evidenced by a decrease in sucrose preference in a sucrose preference test (**Supplementary Fig. 7c,d**). Signals from LHb thus convey strong negative valence to MRN, consistent with previous work linking increased activity within LHb with negative valence, disengagement and depression-like behavior^37–39^. Activation of LHb→MRN input also decreased arousal levels (**Supplementary Fig. 7e-i**). We therefore next tested if LHb→MRN input could also regulate animals’ engagement in different tasks. While suppression of LHb→MRN input did not change the duration of interaction states in the MNOI test, activation of this pathway significantly increased the time mice spent in a disengaged state (**Fig. 5d,e**). LHb→MRN input activation had a similar effect on a learned goal-directed behavior, as it also decreased animals’ engagement in the nose-poke reward test (**Fig. 5f; movie S7**). Activation of the LHb projection to MRN thus has a similar effect on behavior as MRN^SERT^ neuron suppression (**Fig 4b,c**), even though LHb input to MRN is mostly glutamatergic^15,25^.

We thus tested if LHb activation had an excitatory or net inhibitory effect on the activity of serotonergic neurons in MRN, using fiber photometry of calcium signals in MRN^SERT^ neurons combined with optogenetic activation of LHb (**Fig. 5g**). Indeed, LHb activation did not excite MRN^SERT^ neurons (**Fig. 5h**, right), but strongly suppressed their responses to novel objects (**Fig. 5i**). This inhibitory effect is likely mediated by a subset of GABAergic neurons in MRN, since MRN^VGAT^, but not MRN^VGluT2^ neurons, were on average excited by LHb activation (**Fig. 5h**, left and middle). Input from LHb to MRN can thus negatively regulate animals’ engagement levels through inhibition of MRN^SERT^ neurons.

### LHA input to MRN carries positive valence and bidirectionally regulates exploitatory behavior

LHA provides another main input to MRN and has been shown to be important for motivational behaviors^40,41^. We therefore speculated that this input may carry positive valence signals important for sustaining goal-directed behavior. LHA→MRN input is predominantly GABAergic (**Fig. 6b; Supplementary Fig. 8a**), enabling us to manipulate this projection using injection of Cre-dependent ChR2 or eNpHR3.0 into the LHA of VGAT-Cre mice (**Fig. 6b**). Activating GABAergic LHA^VGAT^ axons in MRN significantly increased mice’s preference for the opto-linked port in the self-stimulation test and the opto-linked chamber in the real-time place preference test^25,33^ (**Fig. 6c, Supplementary Fig. 8c,d**), while selectively silencing this pathway had the opposite effect (**Fig. 6c**). Moreover, activation of LHA^VGAT^→MRN input also increased arousal levels (**Supplementary Fig. 8e-i**), and could override animals’ strong aversion to TMT-coated objects, increasing the number of approaches and inducing deep interactions with these aversive objects (**Supplementary Fig. 8j-l**, **movie S7**), similar to the effects of suppressing MRN^VGAT^ neurons. Thus, GABAergic LHA input to MRN conveys strong positive valence while the absence of this input may impose negative valence on animals’ current goal.

We next tested if this projection pathway could also influence exploitatory behavior during interactions with different resources. And indeed, we found that in the MNOI test activation of LHA^VGAT^→MRN input strongly increased the duration of exploitatory states, in which mice showed sustained interactions with one or few objects, and significantly decreased the duration of both exploratory and disengaged states, compared to control mice (**Fig. 6d,e; movie S9**). Moreover, this manipulation increased the time animals spent with each object (**Supplementary Fig. 9a,b**). This finding was corroborated in the T-maze test: after reversal of the reward location, food-restricted mice that had previously been trained to expect a food reward in the now unrewarded T-maze arm continued to prefer this unrewarded arm when LHA^VGAT^**→**MRN input was activated in the central arm of the T-maze, demonstrating a bias towards exploitatory choices (**Fig. 6f**). Notably, these effects closely resembled the increase in exploitatory behavior induced by suppression of MRN^VGAT^ neurons (**Figs. 1d,e,l**).

In contrast, inactivating LHA^VGAT^ axons in MRN in the MNOI test decreased the duration of exploitatory states, while both exploratory and disengaged states were slightly prolonged (**Fig. 6d,e**). Interestingly this manipulation also strongly decreased how long mice spent with individual objects, preventing sustained interactions (**Supplementary Fig. 9c; movie S10**). Accordingly, silencing of LHA^VGAT^ axons in MRN decreased the fraction of exploitatory choices in the T-maze test, causing mice to choose the unrewarded arm more often, even though they were trained to expect reward in the other arm (**Fig. 6g**). Given that LHA^VGAT^ axon silencing did not change animals’ arousal levels (**Supplementary Fig. 8e-i**), and suppressed exploitatory behavior in the MNOI test rather than specifically increasing the time mice spend in an exploratory state, this manipulation likely prevents sustained engagement in a specific action or goal, rather than driving exploratory choices. Interestingly, these effects resembled the results of MRN^VGAT^ neuron activation, which had also disrupted sustained object interactions in the MNOI test (**Supplementary Fig. 9a-c)**. We thus next tested if the influence of LHA^VGAT^ projections in MRN on exploitatory behavior could be mediated through their inhibitory influence on MRN^VGAT^ neurons. And, indeed, fiber photometry recordings of calcium activity in MRN^VGAT^ neurons during optogenetic LHA^VGAT^ neuron manipulation confirmed that activation of LHA^VGAT^→MRN input inhibited MRN^VGAT^ neurons (**Fig. 6h,i**), explaining the similar effect of these neural manipulations on behavior.

Together, our data reveal unexpected brain circuits with a crucial role in regulating how animals interact with environmental resources. Three distinct MRN cell types can differentially drive decisions on whether to exploit current or known options, explore alternative options or disengage from the environment. Suppression of GABAergic neurons in MRN leads to perseverance in current goals and thus biases mice towards exploitatory decisions, by endowing the current, exploitatory option with strong positive valence. One main source of this positive valence are GABAergic neurons in the lateral hypothalamus that inhibit MRN^VGAT^ neurons, and this inhibition is necessary for sustaining current actions or goals. Perseverance in goal-directed actions also necessitates activity of serotonergic MRN neurons that may signal emotionally positive salience. Input to the MRN from the lateral habenula inhibits MRN^SERT^ neurons, likely via a subset of MRN^VGAT^ neurons, and conveys negative valence that decreases task engagement and suppresses sustained interactions. MRN^VGluT2^ neurons appear to operate relatively independently of these pathways, as their activation increases exploratory behavior. Interestingly, suppression of MRN^VGluT2^ neurons had no effect on behavior, indicating that activity in these neurons is not necessary for exploration. However, this pathway provides one route to actively drive exploratory choices. It remains to be determined which upstream brain areas can mediate MRN^VGluT2^ neuron activation, under which circumstances, and whether the induced exploration is goal-directed or random and value-free^42^.

While the MRN is a strong regulator of animals’ choices, it does not act in isolation but as part of complex and distributed subcortical circuits that govern an animal’s motivations and internal states, including dopaminergic systems and the basal ganglia, the interpeduncular nucleus, dorsal raphe nucleus and others^11,13,43–47^. Median and dorsal raphe nuclei are reciprocally connected^17,18,48,49^, and exhibit complementary serotonergic projection patterns, targeting distinct brain areas^48,50^. This suggests a cooperative role of dorsal and median raphe nuclei in balancing interaction states^13,23^. Neocortical input to MRN originates predominantly from prefrontal areas such as anterior cingulate and orbitofrontal cortex^15,16^ (**Supplementary Fig. 10**), areas important for evaluation of expected costs and outcomes of different choices, and updating of value estimates while taking into account outcome uncertainty and other factors^3,4,7,8^. Prefrontal cortex inputs to MRN could exert cognitive control over MRN circuits, eliciting exploratory or exploitatory choices according to higher-order cost-value calculations. In particular, anterior cingulate cortex input to MRN may induce behavioral switches towards exploration through activation of MRN^VGluT2^ neurons^7,51^. How the different MRN cell types interact with each other, and the down-stream circuit mechanisms of the MRN’s influence on exploitatory and exploratory choices remain to be elucidated, however, all three MRN cell types have long-range projections outside the MRN^17,52^.

In summary, MRN can drive either exploratory or exploitatory choices, or disengagement through the differential actions of its three main cell types. Our findings establish MRN as a crucial hub for decision-making and behavioral flexibility. MRN circuits thus may also play an important role in the etiology and possible treatment of major mental pathologies such as depressive or obsessive-compulsive disorders.

## Methods

### Animals and ethics

Mice were housed in individually ventilated cages (IVC) under controlled climate (temperature: 20-24 °C; humidity: 45-65%) in a normal light/dark cycle (12 hr / 12 hr) with ad libitum access to laboratory food pellets and water. Wild-type C57BL/6J (Charles River) mice, and SERT-Cre (Stock #014554, Jackson), VGAT-IRES-Cre (Stock #028862, Jackson) and VGLUT2-IRES-Cre (Stock #016963, Jackson) mice of 8– 12 weeks of age from either sex were used for the experiments. We detected no influence of sex on the results, and data from male and female mice were pooled. All experimental procedures were performed in accordance with UK Home Office regulations (Animal Welfare Act of 2006), under project license PPL PD867676F, following local ethical approval by the Sainsbury Wellcome Centre Animal Welfare Ethical Review Body. Reporting has followed ARRIVE guidelines.

### Virus vector injection and optic fiber implantation

Prior to surgery, mice were subcutaneously injected with the analgesic Metacam (1 mg per kg). Mice were anesthetized with isoflurane (5% induction, 1.2–1.8% maintenance) in oxygen (0.9 L per min flow rate). Body temperature was maintained at 36.5 ℃, using a controlled heating pad. The eyes were protected from light by aluminum foil and from drying by Xailin lubricating eye ointment. Using ear bars, mice were head fixed on a stereotactic device (Kopf, model 940) and using a scalpel blade the scalp was cut along with the midline to expose the skull. Small craniotomies were made with a dental drill (0.4 or 0.5 mm diameter) and AAV virus (60 nl) was injected into the target brain regions using a pulled glass pipette (∼ 20 µm inner diameter at the tip) and a programmable nanoliter volume injector (Nanoject III, Drummond Scientific). Fifteen minutes after the injection, the glass pipette was retracted and the incision was either sutured or glued (Vetbond), or optic fibers (for optogenetics: 200 µm diameter; for photometry: 400 µm diameter) were implanted. After cementing a custom-designed metal head plate to the skull (using light-cure Tetric EvoFlow cement (Ivoclar Vivadent), empowered by OptiBond Universal (Kerr) primer), the optic fibers were inserted 200 µm (for optogenetics) or 50 µm (for photometry) above the target brain regions and cemented to the skull. After recovery, the animals were returned to their cage. Eighteen to twenty days after the virus injection and fiber implantation surgery, mice with expression of an optogenetic opsin (ChR2 for neuronal or axonal activation, ChrimsonR for neuronal activation during photometry recording, stGtACR2 for neuronal suppression, or eNpHR3.0 for axonal suppression), tdTomato (control for optogenetic stimulation) or GCaMP6f (for calcium photometry recording) were used for optogenetics/photometry experiments. After recovery, the animals were returned to their cage.

Brain regions and coordinates (from Bregma) used for virus injections: MRN (AP: −4.4 mm, ML: 0.0 mm, DV: 4.3 mm or AP: −5.5 mm, ML: 0.0 mm, DV: 4.4 mm, AP angle: 14 deg.), LHb (AP: −1.8 mm, ML: 0.4 mm, DV: 2.7 mm) and LHA (AP: −5.2 mm, ML: 1.0 mm, DV: 5.2 mm). For optogenetic stimulation, we used AAV9-hEF1a-DIO-mCherry-hChR2 (University of Zurich; V80-9, a gift from Karl Deisseroth), AAV1-CAG-hChR2-tdTomato (virus made in the host institute, using Addgene plasmid #28017 a gift from Karel Svoboda), AAV9-hSyn-SIO-FusionRed-stGtACR2 (virus made in the host institute, using Addgene plasmid #105677, a gift from Ofer Yizhar), AAV1-Ef1a-DIO-eNpHR3.0-EYFP (Addgene #26966-AAV1, a gift from Karl Deisseroth), AAV1-hSyn-eNpHR3.0-EYFP (virus made in the host institute, using Addgene plasmid #26972, a gift from Karl Deisseroth), AAV1-hSyn-DIO-ChrimsonR-tdTomato (virus made in the host institute), AAV1.Syn.ChrimsonR.tdTomato (Addgene # 59171-AAV1, a gift from Edward Boyden) and AAV1-CAG-tdTomato (Addgene #59462-AAV1, a gift from Edward Boyden), for fiber photometry recordings, AAV1-Syn-flex-GCaMP6f (virus made in the host institute, using Addgene plasmid #100833, a gift from Douglas Kim & GENIE Project) and for retrograde tracing experiments, retroAAV-CAG-GFP (virus made in the host institute, using Addgene plasmid #37825, a gift from Edward Boyden). Viruses for optogenetics, photometry and retrograde tracing were diluted to ∼ 5 × 10^12^ vg/ml, ∼ 2 × 10^12^ vg/ml and ∼ 8 × 10^12^ vg/ml, respectively.

### Optogenetics

Laser stimulation protocols were created and run through custom-made scripts in MATLAB or Python. For neuronal or axonal activation by ChR2 and neuronal suppression by stGtACR2 a 473 nm laser (OBIS 473nm LX 75mW LASER SYSTEM, Coherent), coupled to a 200 µm fiber patch cable through an achromatic fiber port (Thorlabs), was used. For axonal suppression by eNpHR3.0 and neuronal activation by ChrimsonR we used 594 nm (OBIS 594nm LS 40mW LASER SYSTEM: FIBER PIGTAIL: FC, Coherent) and 647 nm (OBIS 647nm LX 120mW LASER SYSTEM, Coherent; coupled to a 200 µm fiber patch cable through an achromatic fiber port (Thorlabs)) lasers, respectively. The laser frequency was 20 Hz (with 50% duty-cycle pulses) for depolarizing opsins (ChR2 and ChrimsonR) and 0 Hz for hyperpolarizing opsins (stGtACR2 and eNpHR3.0). The peak laser power at the tip of the fiber was ∼2 mW. Using a Pulse Pal pulse generator (Open Ephys), each laser pulse for stimulating stGtACR2 was initiated and followed by a linear ramp-up and ramp-down of 500 ms, respectively. Stimulation of other opsins was done by square pulses.

### Multi novel object interaction test

Mice were habituated to the experimenters and the experimental box (40 cm × 40 cm × 50 cm) every day for three weeks. For the last three days before the experiment, mice were also habituated to one novel object in the box to minimize stress in response to novel objects. Multi novel object interaction tests were designed in a free access manner. Mice were put in the box for 15 minutes before the experiment. Then 20 novel objects (with different shapes, colors, textures and materials) were put in random locations, and mice were allowed to interact with the objects for 2 min with optogenetic laser stimulation throughout the test duration (alternating periods of laser on 4.7 s and laser off 0.3 s). All objects were small (length between 0.5-1.0 cm) and light enough for the mice to be able to pick up and displace. Behavior was video recorded using one camera from the top, fixed to the ceiling of the box, and two side cameras (Raspberry Pi cameras (V2 module) with a frame rate of 25 fps).

Videos were labeled frame by frame using JAABA^1^. Analyzers were blind to the experimental groups of mice. Behaviors displayed during object interaction were categorized as approaching: turning of the head towards the object accompanied by a body movement decreasing the distance between the mouse and the object; approaches ended when the mouse was at 0.5cm distance to the object: sniffing: location within whisking distance of the object (closer than 0.5 cm) and facing and seemingly ‘focusing on’ the object for at least 5 frames, this can include touching the object with the snout, but not biting it; deep interaction: at least one of the following actions are taken: biting (taking hold of the object between its jaws and/or making nibbling motions with its head, but not walking around with the object), grabbing (holding of the object between the front paws, or standing over the object and blocking the object, effectively preventing it from moving away), and/or carrying (holding the object in its mouth and simultaneously walking around the box with it, effectively displacing the object); leaving the object: moving away from an object after sniffing or deep interaction (until the nose is turned 0.5 cm away from the object). Each of these actions was attributed to one of the 20 objects. JAABA labels were imported to MATLAB for further analysis. After extracting mouse positions and movement speed, using custom-made scripts in MATLAB, times that the mouse was not performing the above actions (approaching, sniffing, deep interaction and leaving) were attributed to sitting or walking (binarizing at movement speed of 0.05 cm/s).

### Hidden Markov model

We used a hidden Markov model (HMM) to extract states that may underlie different sequences of actions. The HMM requires an estimated transition matrix, an estimated emission matrix and a sequence as input. The sequence was generated from the data of assigned mouse behaviors during the multi object interaction test. We created a non-overlapping sequence vector, with just one action happening at each time. This was achieved by creating a ranking sequence in which non-object-interacting behavior (sitting and walking) was the lowest, then leaving, approaching, sniffing and deep interacting was the highest rank, and the highest rank was chosen as current action. We assumed that the higher the rank was, the more important information it conveyed of the mice’s state for investigatory behavior. We set the model to three states, because we hypothesized three underlying states important for object interactions, an exploitatory state in which mice persevere in interaction with the current object, an exploratory state in which mice switch between multiple objects with short interactions, and a disengaged state without object interactions. Since we did not have a priori information on the transitions between the states, our starting estimate for the transition matrix contained equal probabilities for all transitions, but starting with a random transition matrix did not change the results. We generated the initial estimated emission matrix by predicting the likelihood of a behavior present in a certain state. Using a Baum-Welch algorithm^2^, we trained the model on control mice to estimate the transition and emission probabilities for the HMM (**Fig. 1b and Supplementary Fig. 1a**). Setting the model to four states yielded similar results, i.e. an exploitatory state (mainly consisting of deep interaction), an exploratory state (mainly consisting of sniffing, approaching and leaving of multiple objects, with probabilities of 0.71, 0.16 and 0.11, respectively), and two states without object interactions. As this test was focused on object interactions, we thus chose to use the three-state HMM. For all other groups of mice (including mice used for optogenetic and photometry experiments), the states’ probabilities at each time bin (bin size 0.5 seconds) were decoded from their vectors of rank-number actions based on the emission and transition matrices trained on control mice. Finally, each time bin was attributed to the state with maximum likelihood, resulting in a sequence of states. We used the built-in HMM functions hmmtrain and hmmdecode in MATLAB to acquire and evaluate the transition and emission matrices for control mice and estimate the states’ probabilities for other mice. To estimate the conditional transition probability between different states in **Supplementary Fig. 1b**, one time bin was used for the duration of each state, such that transition probabilities were only calculated for time points of state changes.

### Real-time place preference test

Mice with expression of an optogenetic opsin (ChR2, stGtACR2 or eNpHR3.0) or tdTomato (control) underwent a real-time place preference test (RTPPT) in a custom-made two-chamber acrylic box (60 cm × 30 cm × 30 cm (l × w × h)). After 10 minutes of habituation to the box, one of the two chambers was paired with laser stimulation (triggered by entering the chamber). The laser-coupled chamber was randomly assigned. The total test duration was 10 minutes. Mouse behavior was analyzed using Bonsai software (https://bonsai-rx.org/). Using a custom-made MATLAB script, the preference for the opto-linked chamber was calculated as follows: 100% × (duration of time spent in the opto-linked chamber - duration of time spent in the non-stimulation chamber) / total time).

### Self-stimulation test

Mice with expression of an optogenetic opsin (ChR2, stGtACR2 or eNpHR3.0) or tdTomato (control) were habituated for 1 hour to a custom-made experimental box (40 cm × 40 cm × 50 cm; l × w × h) with a two-port nose-poke system one day before the test. During the habituation period, drops of 10% sucrose water were delivered through both ports to habituate the mice to the nose-poke ports. No sucrose water was delivered during the test. Instead, an infrared sensor in only one nose-poke port triggered optogenetic stimulation when the mouse entered the port with its nose, and stimulation continued throughout the time-period the mouse’s nose remained in the nose-poke port. An infrared sensor in the other nose-poke port did not trigger any stimulation. The test lasted 1 hour. The number of returns to each port was detected by the sensors (connected to an Arduino UNO microcontroller board) and preference for entering the opto-linked nose-poke port was calculated as follows: 100% × (number entries into the opto-linked nose-poke port - number of entries into the non-stimulation nose-poke port)/(number of entries into the opto-linked nose-poke port + number of entries into the non-stimulation nose-poke port).

### T-maze tests

To measure the effects of optogenetic stimulation on exploratory vs exploitatory choices based on acquired choice-outcome knowledge, we trained food-restricted mice on a T-maze task, in which a food reward was provided consistently only in one specific arm and not the other. Training started after two days of 15-minute habituation to the T-maze, to the experimenter and to a soft towel by which they were placed in and picked up from the T-maze. Each training trial started with placement of the mouse at the start of the central arm and ended with the mouse turning into either of the choice arms (left or right). At the end of each trial the entrance of the choice arms was blocked to prevent the mouse from returning to the other arms. After reaching the end of the non-rewarded arm or after eating the food reward at the end of the rewarded arm, mice were gently picked up in the soft towel and placed back at the start of the central arm. After mice achieved more than 90% correct trials in two sequential 50-trial sessions (well-trained; mean ± std: 7.42 ± 2.66 sessions), they entered one of the two tests. In the exploration test (55 trials), the food reward was provided in the same arm as during training, and optogenetic stimulation was delivered in the central arm of the T-maze until after mice chose one of the arms. The percentage of trials in which they entered the non-rewarded arm indicated their tendency to explore. In the exploitation test (55 trials) the reward location in the T-maze was reversed, i.e. the previously rewarded arm was not rewarded anymore. In this scenario, normally mice start exploring the other, previously non-rewarded arm and eventually learn to expect reward only at this location. Mice in this reversal-learning test received optogenetic stimulation in the central arm of the T-maze until after they chose one of the arms. The percentage of trials in which they entered the previously rewarded arm (i.e. the currently non-rewarded arm) indicated their tendency to exploit.

### Nose poke-reward association task

To measure the effect of optogenetic stimulation on task engagement, we used a previously described nose poke-reward association task^3^. After three days of water restriction, mice with expression of an optogenetic opsin (ChR2, stGtACR2 or eNpHR3.0) or tdTomato (control) were trained to enter their nose into a nose-poke port on one wall of a custom-made operant chamber (25 cm × 20 cm × 30 cm (l × w × h); in a sound-attenuating box) in order to receive a water reward from a lick spout on the wall opposite to the nose-poke port. At the start of each trial, the nose-poke port was illuminated. Upon completion of a successful nose-poke, the light in the nose-poke port was turned off, and a white noise sound was turned on to indicate the availability of reward. A water reward was delivered to the lick spout, via a solenoid valve, upon licking. A nose-poke entry followed by a lick was considered a single completed trial. Mice were free to run back and forth between the nose-poke port and the reward spout and complete trials at their own pace. Training continued daily until mice were able to complete more than 85 trials per session in two sequential daily 30-minute sessions (mean ± std: 10.40 ± 4.35 sessions). Following training, mice performed the same task during which they received 3 epochs of 2-minute optogenetic stimulation with random off-stimulation time intervals (not less than 2 minutes) between 2 min and 20 min from the session’s onset. The number of completed trials indicated the engagement level. Data acquisition and stimulation was performed using a Pycontrol state-machine (https://pycontrol.readthedocs.io/en/latest/).

### Nose poke-reward association task with distractor nose-poke ports

To measure the effect of optogenetic stimulation on evoking random exploration, we modified the nose poke-reward association task, by adding two nose-poke ports without function at the side walls. Following training in the standard nose poke-reward association task, mice with expression of ChR2 in VGluT2+ or VGAT+ neurons in MRN, or tdTomato (control) mice were tested in the presence of the two additional nose-poke ports, used as distractors to motivate exploration. Only an entry into the previously learned nose-poke port followed by a lick from the lick spout resulted in water-reward delivery. At the beginning of the test, mice were given a 5-minute period to explore all the nose-poke ports and understand that the additional nose-poke ports are not associated with reward. During the following 20 minutes, mice received optogenetic stimulation (for a random time period between 1 and 2 seconds) in 25-35% of trials upon entering into a 0.5 cm radius around the reward-associated nose-poke port. From laser onset until completion of the trial (receiving a water-reward), the number of times mice interacted with the distractor nose-poke ports or the lick spout (without initiating the trial) was counted as number of interactions with distractors in this trial. Data acquisition and stimulation was performed using a Pycontrol state-machine and data analysis was performed by a custom-made MATLAB script.

### TMT aversion test

Mice were habituated to the experimenter for three days, 30 minutes a day, and to the experimental box (40 cm × 40 cm × 50 cm (l × w × h)) for 30 minutes the day prior to the experiment. For the test, mice were habituated in the box for 15 minutes, after which a small object covered with 3 µl TMT (BioSRQ: purity > 90.0%) was placed inside the box for 2 minutes. Optogenetic stimulation was applied for the 2-minute duration, with repeated pulse trains of 4.7 s laser on and 0.3 s laser off (to avoid overstimulation of neurons and tissue damage by heat accumulation). Mouse behavior was video recorded (Raspberry Pi camera (V2 module); with 25 fps) and videos were labeled frame by frame using JAABA^1^. Analyzers were blind to the experimental groups of mice. Behaviors displayed during the test were categorized as: approaching (approaching towards the object, from start of body movement until reaching proximity of 0.5 cm), interacting (sniffing, grabbing, carrying or biting the object) and retreating (moving away, upon reaching the object, in the opposite direction to the approach). JAABA labels were imported to MATLAB for further analysis. In addition, mouse position and movement speed were extracted using custom-made MATLAB scripts. A retreat upon reaching the TMT-object was counted as an escape if the average speed of the retreat exceeded 0.5 cm/s within the first second. Varying the speed threshold (0.3, 0.4, 0.5 or 0.6 cm/s) did not influence the results. Escape probability was calculated as the number of approaches leading to escape divided by the total number of approaches.

### Visually-evoked fear response test

Mice with expression of ChR2 in VGLUT2+ MRN neurons underwent a visually-evoked fear response test to see whether activation of these neurons changes their fear response to an innately threatening visual stimulus, an overhead expanding, i.e. looming, black disc^4^. Experiments assessing escape behavior in response to looming stimuli were performed in a custom-made transparent acrylic arena (80 cm × 26 cm × 40 cm (l × w × h)) with a red-tinted acrylic shelter (14 cm × 14 cm × 14 cm (l × w × h)) placed on one end (safe zone) and overhead looming stimuli presented on the other end (threat zone)^5,6^. The arena was placed in a large light-proofed and sound-attenuating box with a near-IR GigE camera (acA1300-60 gmNIR, Basler; with a frame rate of 50 fps) fixed on the ceiling to video record the behavior. To display looming stimuli, a projector (HF85JA, LG) was mounted in the box and back-projected via a mirror onto a suspended horizontal screen (60 cm above arena floor, 100 cm × 80 cm; ‘100 micron drafting film’, Elmstock). The screen was kept at a constant background luminance level of 9 cd × m−2. The arena was illuminated by four infrared LEDs to ease tracking of the mice. Video recording and optogenetics laser stimulations were triggered through Bonsai (https://bonsai-rx.org/). Mice were placed in the arena 20 minutes before the test to habituate to the new environment. Stimulation was triggered manually by a keyboard when the mouse reached the threat zone. The stimulus was only triggered when the mouse was facing and walking toward the threat zone, with an inter-stimulus interval of at least two minutes. Each visual stimulation consisted of three consecutive looming stimuli (expanding black spot at a linear rate of 55 deg/s) in a 3-second period. Visual stimuli of 10%, 50% and 90% contrasts were presented in a randomized order. Laser and non-laser trials of the same stimulus contrast were always presented as paired trials, in a randomized but consecutive order (10 repetitions × 3 contrasts × 2 laser conditions). Optogenetic stimulation in laser trials started 0.5 seconds before the visual stimulus. Position and running speed of the center of the mouse were processed using a custom-made MATLAB script. A successful escape was defined as a return to the shelter with an average running speed higher than 0.4 cm/s (varying this threshold to 0.3 or 0.5 cm/s did not change the significance of the results) in the two seconds after the stimulus onset and reaching the shelter within five seconds of stimulus onset.

### Sucrose preference test

Sucrose preference test (SPT) is frequently used to measure anhedonia in mice based on a two-bottle choice paradigm^7,8^. A decreased responsiveness to rewarding sucrose compared to a control group indicates anhedonia. On the first day of habituation, mice with expression of ChR2 in LHb input to MRN and control mice were provided with two identical bottles of water and on the second day with two identical bottles of 1% sucrose water for 24 hours, respectively. During the next 4 days, mice were provided with two bottles of water, while they were receiving repeated optogenetic stimulation for 24 hours (60 s laser on, every 240 s). The next day, after 12 hours of water deprivation, mice were tested with one bottle of water and one bottle of 1% sucrose water (for 12 hours), without optogenetic stimulation. Sucrose preference was calculated by the percentage of sucrose water consumed relative to the total liquid consumption.

### Arousal level measures and analysis

For these experiments, mice were habituated for three days (30 min a day) to be head-fixed on a running-wheel in a sound-attenuating box with dim ambient light. Mice with expression of an optogenetics opsin (ChR2, stGtACR2 or eNpHR3.0) or tdTomato (controls) were used for measuring changes in physiological arousal level caused by optogenetic stimulation. Mice were head-fixed and an infrared LED light was directed to the face to illuminate the pupil (mounted 50 cm away). Mice received optogenetic stimulation for 3-second periods (30 trials with 12-second intervals). The effect of optogenetic stimulation on the arousal level was monitored by recording videos of the pupil and whiskers (using two cameras (22BUC03, ImagingSource) with frame rates of 30 fps). Whisker activity during each frame was calculated as the absolute difference between the color intensity of each pixel of this frame and the previous frame, averaged over all pixels. Z-scored pupil size and whisker activity were determined using custom-made scripts in MATLAB.

### Calcium photometry recording and data analysis

Mice were habituated to the experimenters for three days (30 min a day) and to the experimental box (40 cm × 40 cm × 40 cm) for more three days (15 min a day). Three weeks after the GCaMP6 viral injection and the optic fiber implantation, calcium activity of MRN neurons was recorded in freely moving mice. Recording from SERT-Cre mice was performed in three different conditions, in the presence of multiple novel objects, of food pellets, or of TMT objects. Recordings from VGAT-Cre and VGLUT2-Cre mice were done in presence of multiple novel objects. Movies of mouse behavior and fluorescence were recorded simultaneously using Raspberry Pi cameras (V2 module, frame rate: 25 fps) and a fiber photometry system (optics from Doric Lenses; acquisition board and recording software from PyPhotometry (https://pyphotometry.readthedocs.io/en/latest); sampling rate: 130 Hz), respectively. Excitation lights at the tip of the optic fiber were adjusted to around 30 µW. The data for fiber photometry was analyzed using a custom-made MATLAB program. A linear drift correction was applied to raw signals of calcium-dependent (GCaMP excited at 473 nm) and isosbestic fluorescence (GCaMP excited with 405 nm light) to correct for slow changes such as photobleaching. To correct for non-calcium dependent signals and artifacts, the isosbestic fluorescence trace was linearly fitted to the calcium-dependent GCaMP fluorescent signal and subtracted, providing a measure of relative transient changes in fluorescence (dF/F). Mean baseline was taken as the average dF/F signal of the entire recording session. Subsequently, z-scored dF/F was calculated by subtracting the mean baseline and dividing by the standard deviation of the baseline distribution. Behaviors were analyzed using JAABA and MATLAB, similar to experiments with optogenetic stimulation. Data points in **Figs. 1g and 2e** are z-scored dF/F values averaged over 3 seconds after the onset of behaviors stated in the figure legends. Data points in bar graphs in **Figs. 1h and 2f** are z-scored dF/F values averaged over the time course of the associated interaction states.

### Histology and microscopy

For determining inputs and outputs of specific brain areas, and for histological confirmation of injection sites and fiber locations, at the end of experiments mice were euthanized by an overdose of pentobarbital (intraperitoneal injection, 80 mg × kg−1) and transcardially perfused (0.01 M phosphate buffered saline (PBS), followed by 4% paraformaldehyde (PFA) in PBS). After extraction, brains were post-fixed by 4% PFA solution overnight and consequently embedded in 5% agarose (A9539, Sigma). Imaging was performed using a custom-built automated serial-section two-photon microscope (Mayerich et al., 2008; Ragan et al., 2012). Coronal slices were cut at a thickness of 40 μm, and images were acquired from 2-8 optical planes (every 5-20 μm) with ∼2.3 μm/pixel resolution. Scanning and image acquisition were controlled by ScanImage v5.5 (Vidrio Technologies, USA, Pologruto et al., 2003) using a custom software wrapper for defining the imaging parameters (https://zenodo.org/record/3631609).

### Overall experimental design and analysis

No statistical methods were used to predetermine sample sizes, but our sample sizes were determined based on previous studies^9,10^. The order of animals in different experimental groups was randomly assigned. Experimenters were not blind to the experimental conditions, but the collected data were encoded blindly and analyzers were blind to experimental condition. All data were analyzed using JAABA, MATLAB, Python and Bonsai.

### Statistical analysis

Data are represented as median ± bootstrap standard error, unless stated otherwise. All statistical analyses were performed using InVivoTools MATLAB toolbox^11–13^ (https://github.com/heimel/InVivoTools) or custom-made MATLAB scripts. First, normality of the data distribution was tested, using a Shapiro-Wilk normality test. To assess group statistical significance, if data were normally distributed we used parametric tests (i.e. t-test and paired t-test for non-paired and paired comparisons, respectively), and otherwise non-parametric tests (i.e. Mann-Whitney U test and Wilcoxon signed rank test for non-paired and paired comparisons, respectively), followed by a Bonferroni p-value correction for multi group comparisons. For multi group comparisons with subgroups within groups we used nested one-way ANOVA. To statistically compare distributions of discrete data across different groups, chi-square test was used. Individual data points are shown in the figures. All statistics used in figures and supplementary figures are listed in the figure legends.

## Acknowledgements.

We thank Michelle Li for animal husbandry and genotyping; Rob Campbell for help with microscopy; Anna Seggewisse for help with animal training; the S.B.H. laboratory and the Thomas Mrsic-Flogel laboratory for the discussions; Thomas Mrsic-Flogel and Tim Behrens for feedback on the manuscript and discussions; This work was supported by the Sainsbury Wellcome Centre Core Grant from the Gatsby Charitable Foundation and Wellcome (S.B.H., GAT3755 and 219627/Z/19/Z) and an European Research Council Starting Grant (S.B.H., HigherVision 337797).

## Author contributions

M.A. and S.B.H. conceived the study. M.A. set up the experiments and analysis pipelines, performed the experiments and analyzed the data. M.Y.S. and P.Z. assisted with histology. M.Y.S. assisted with surgical procedures. M.Y.S., P.Z. and I.L.M.R. conducted animal habituation, training, and behavior analysis, and assisted with optogenetic and photometry experiments. J.D. assisted with setting up the experiments.

## Data and code availability

Data analysis codes of this study are available from the corresponding authors upon request.

