## Supplemental Materials for "A subcortical switchboard for exploratory, exploitatory, and disengaged states"

### Supplementary Figures

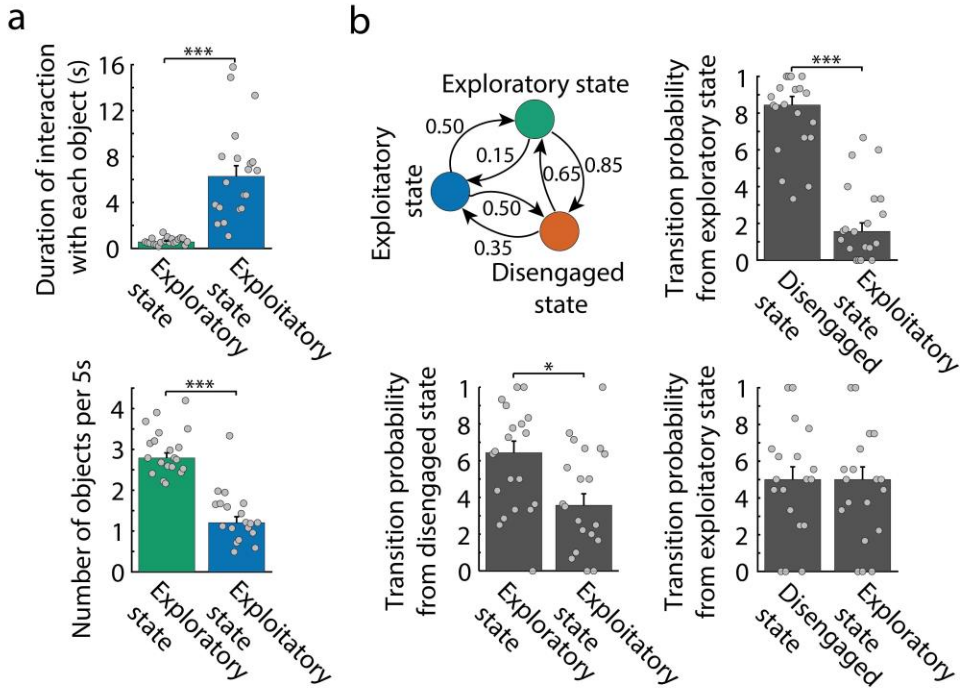


**Supplementary Figure 1. Interaction states extracted by hidden Markov model and transition probabilities between the states**

**a,** Average duration of object interactions (top, $P = 2.3\times10^{-6}$, two-sided paired t-test) and frequency of object interaction (bottom, P = 0.0001, Wilcoxon signed rank test) in exploratory (green) and exploitatory (blue) states in control mice during the MNOI test. **b,** Median transition probabilities at the time of switching between the three interaction states in control mice. Bar graphs show the transition probability from each state to the other two states (median values and individual experiments). From exploratory state to disengaged vs. exploitatory state: P = 0.0003, Wilcoxon signed rank test; from disengaged state to exploratory vs. exploitatory state: P = 0.0302, two-sided paired t-test; from exploitatory state to disengaged state vs. exploratory state: P = 0.8047, two-sided paired t-test, N = 20 experiments from 10 mice). *: p-value < 0.05, ***: p-value < 0.001. Bars depict median, error bars are bootstrapped standard error and circles indicate individual experiments.

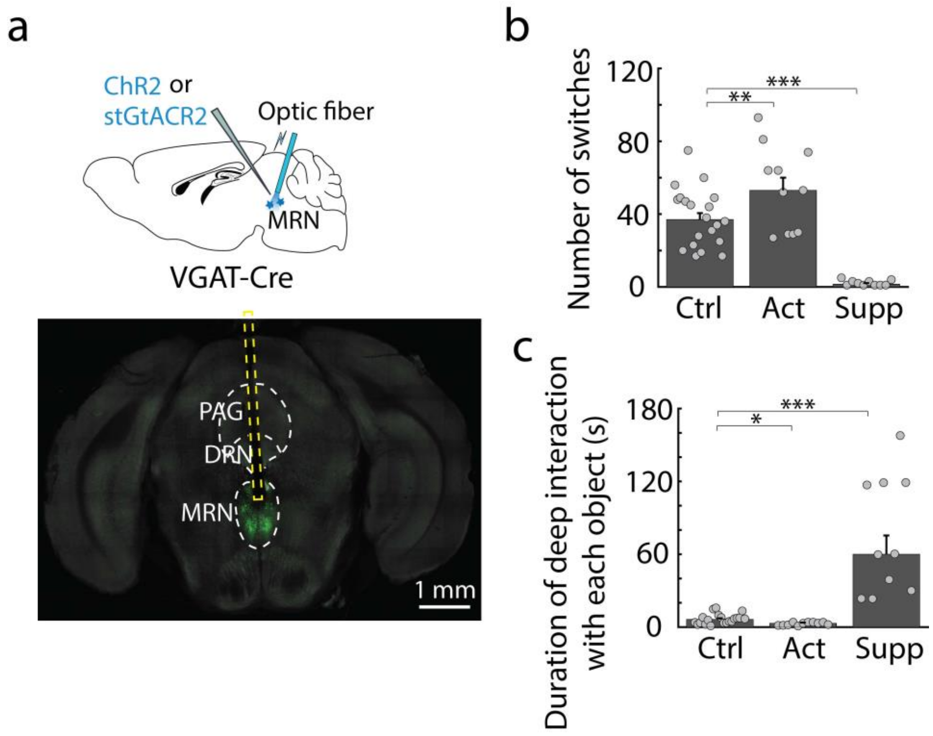


**Supplementary Figure 2. VGAT+ MRN neurons bidirectionally modulate how often mice switch between objects and how long they interact with each object**

**a,** Schematic of experimental design: optogenetic activation or suppression of VGAT+ MRN neurons, using ChR2 or stGtACR2 (top) and example image of virus expression in VGAT+ MRN neurons with optic fiber position. DRN: dorsal raphe nucleus, MRN: median raphe nucleus, PAG: periaqueductal gray. **b,** Number of switches between objects during the MNOI test in control (ctrl) mice and mice with activation (act) and suppression (supp) of VGAT+ MRN neurons (ctrl vs. act: $P = 0.00591$, ctrl vs. supp: $P = 2.3\times10^{-7}$, two-sided t-test with Bonferroni multi-comparison correction). N = 20, 11 and 10 experiments from 10, 6 and 5 mice in control, VGAT+ activation and VGAT+ suppression groups. **c,** Duration of deep interactions (when mice grab, bite or carry the object) with each object during the MNOI test of mice in **b** (ctrl vs. act: $P = 0.0102$, ctrl vs. supp: $P = 1.6\times10^{-6}$, two-sided t-test with Bonferroni multi-comparison correction). *: p-value < 0.05, **: p-value < 0.01, ***: p-value < 0.001.

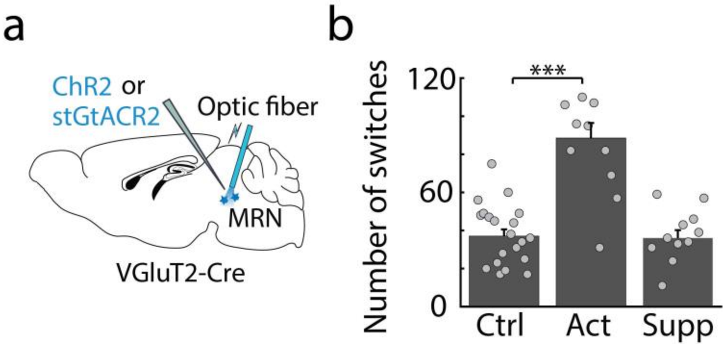


**Supplementary Figure 3. Activation of VGluT2+ MRN neurons increases the number of switches between objects**

**a,** Schematic of experimental design: optogenetic activation or suppression of VGluT2+ MRN neurons, using ChR2 or stGtACR2. **b,** Number of switches between objects during the MNOI test in control mice (ctrl) and mice with activation (act) and suppression (supp) of VGluT2+ MRN neurons (ctrl vs. act: $P = 3.2\times10^{-6}$, ctrl vs. supp: $P > 0.9999$, two-sided t-test with Bonferroni multi-comparison correction). N = 20, 10 and 11 experiments from 10, 5 and 5 mice in control, VGluT2+ activation and VGluT2+ suppression groups. ***: p-value < 0.001. Bars depict median, error bars are bootstrapped standard error and circles indicate individual experiments.

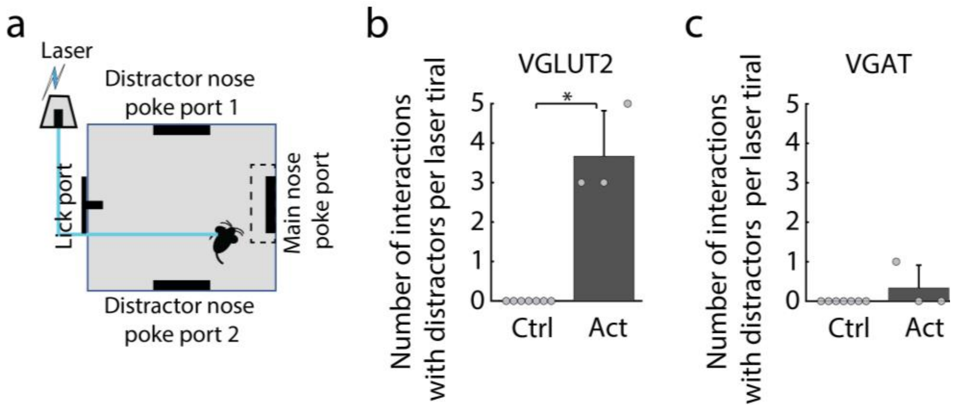


**Supplementary Figure 4. Activation of VGluT2+ but not VGAT+ MRN neurons actively drives exploratory behavior**

**a,** Schematic of nose-poke reward association task with distractor nose poke ports, which have no association with reward. Mice with expression of ChR2 in VgluT2+ or VGAT+ MRN receive brief laser stimulation (1-2 s) upon entering a region of interest around the reward-associated (main) nose poke port in a subset of trials (see Methods). **b,** Median number of interactions with distractors per laser stimulation trial over all the completed laser stimulation trials in control mice and mice with optogenetic activation of VGluT2+ MRN neurons ($P = 0.0027$, two-sided chi-square test, $N = 7$ vs. 3 mice). **c,** The same as **b**, but in control mice and mice with optogenetic activation of VGAT+ MRN neurons ($P > 0.9999$, two-sided chi-square test, $N = 7$ vs. 3 mice). **: p-value < 0.01. Bars depict median, error bars are bootstrapped standard error and circles indicate individual experiments.

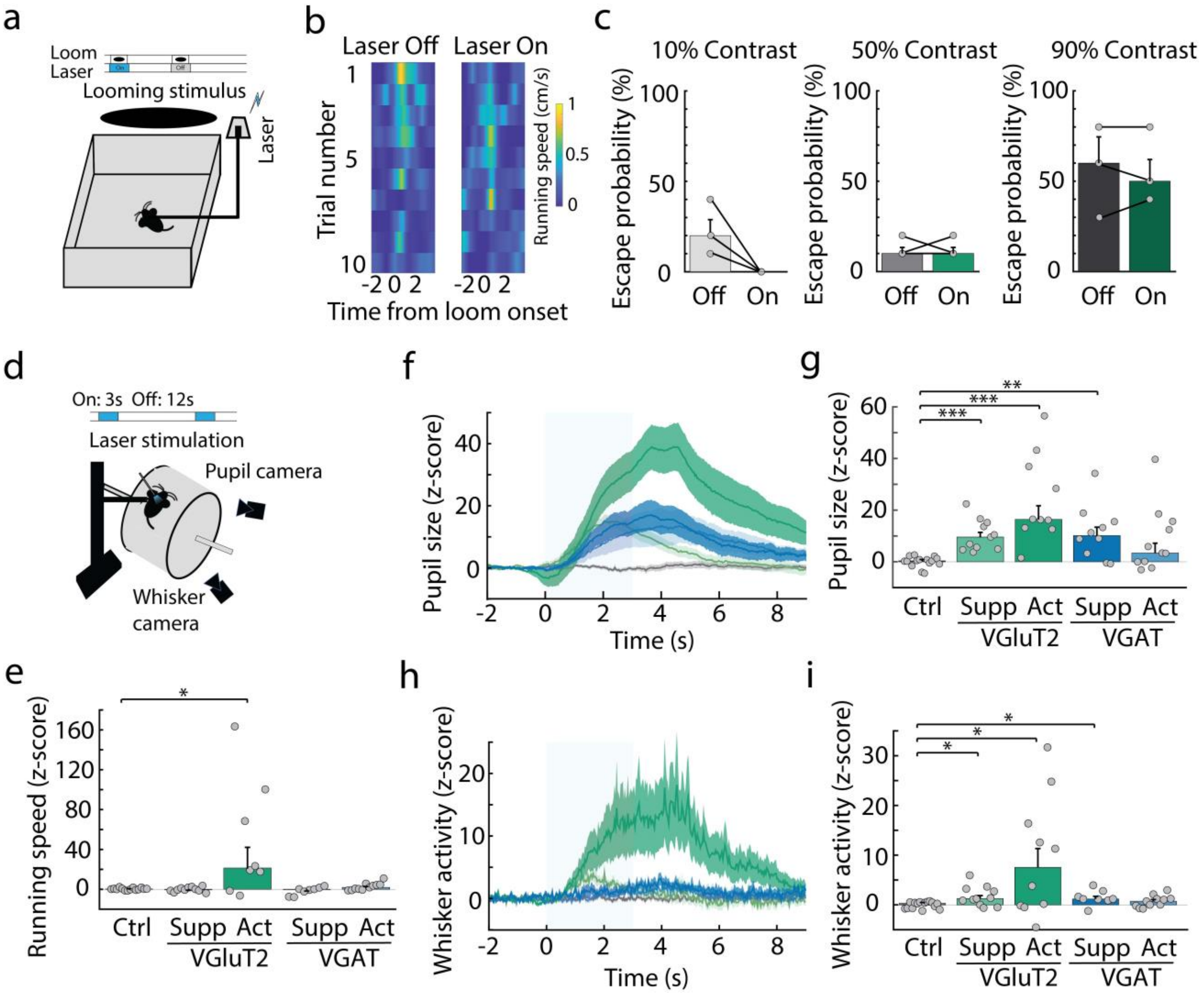


**Supplementary Figure 5. Modulation of arousal and valence by VGluT2+ and VGAT+ MRN neurons**

**a,** Schematic of the experimental design for quantifying fear responses to an innately aversive looming stimulus, with three different stimulus contrast levels (10%, 50%, and 90%), with and without optogenetic activation of VgluT2+ MRN neurons. **b,** Heatmaps show running speed (cm/s) of an example mouse in response to 90% contrast looming stimuli in laser off and laser on trials. **c,** Escape probability of mice in response to looming stimuli with different contrasts, during activation of VgluT2+ MRN neurons (on) and control trials (off). N = 3 mice, 10% contrast: P = 0.1181; 50% contrast: P > 0.9999; 90% contrast: P > 0.9999, two-sided paired t-test. **d,** Schematic of the experimental design to examine the effects of optogenetic manipulation of VGluT2+ and VGAT+ MRN neurons on running speed and arousal level, measured using pupil size and whisker activity. **e,** Z-scored running speed evoked by optogenetic stimulation in control mice and mice with activation or suppression of VGluT2+ or VGAT+ MRN neurons. N = 15, 11, 9, 8 and 11 mice, respectively; P > 0.9999, P = 0.0229, P = 0.3880 and P = 0.1210 for comparing control mice to activation or suppression of VGluT2+ neurons, and activation or suppression of VGAT+ neurons, respectively; two-sided t-test with Bonferroni multi-comparison correction. **f,** Z-scored pupil size over time (mean ± s.e.m.) in control mice (grey) and mice with activation or suppression of VGluT2+ and VGAT+ MRN neurons, averaged over trials and aligned to onset of optogenetic manipulation. Light blue bar indicates the laser stimulation period. **g,** Median z-score of pupil size during laser stimulation from traces in **f**. N = 15, 11, 10, 10 and 11 mice; P =

$8.0\times10^{-6}$, P = $6.7\times10^{-5}$, P = 0.0015 and P = 0.0660 for comparing control mice to activation or suppression of VGluT2+ neurons, and activation or suppression of VGAT+ neurons, respectively; two-sided t-test with Bonferroni multi-comparison correction. **h,i,** same as **f,g** but for z-scored whisker activity (summation of absolute frame-by-frame differences in pixel luminance). N = 15, 11, 10, 8 and 11 mice, respectively; P = 0.0165, P = 0.0195, P = 0.0153 and P = 0.2356 for comparing control mice to activation or suppression of VGluT2+ neurons, and activation or suppression of VGAT+ neurons, respectively; two-sided t-test with Bonferroni multi-comparison correction. *: p-value < 0.05, **: p-value < 0.01, ***: p-value < 0.001. Bars depict median, error bars are bootstrapped standard error and circles indicate individual experiments.

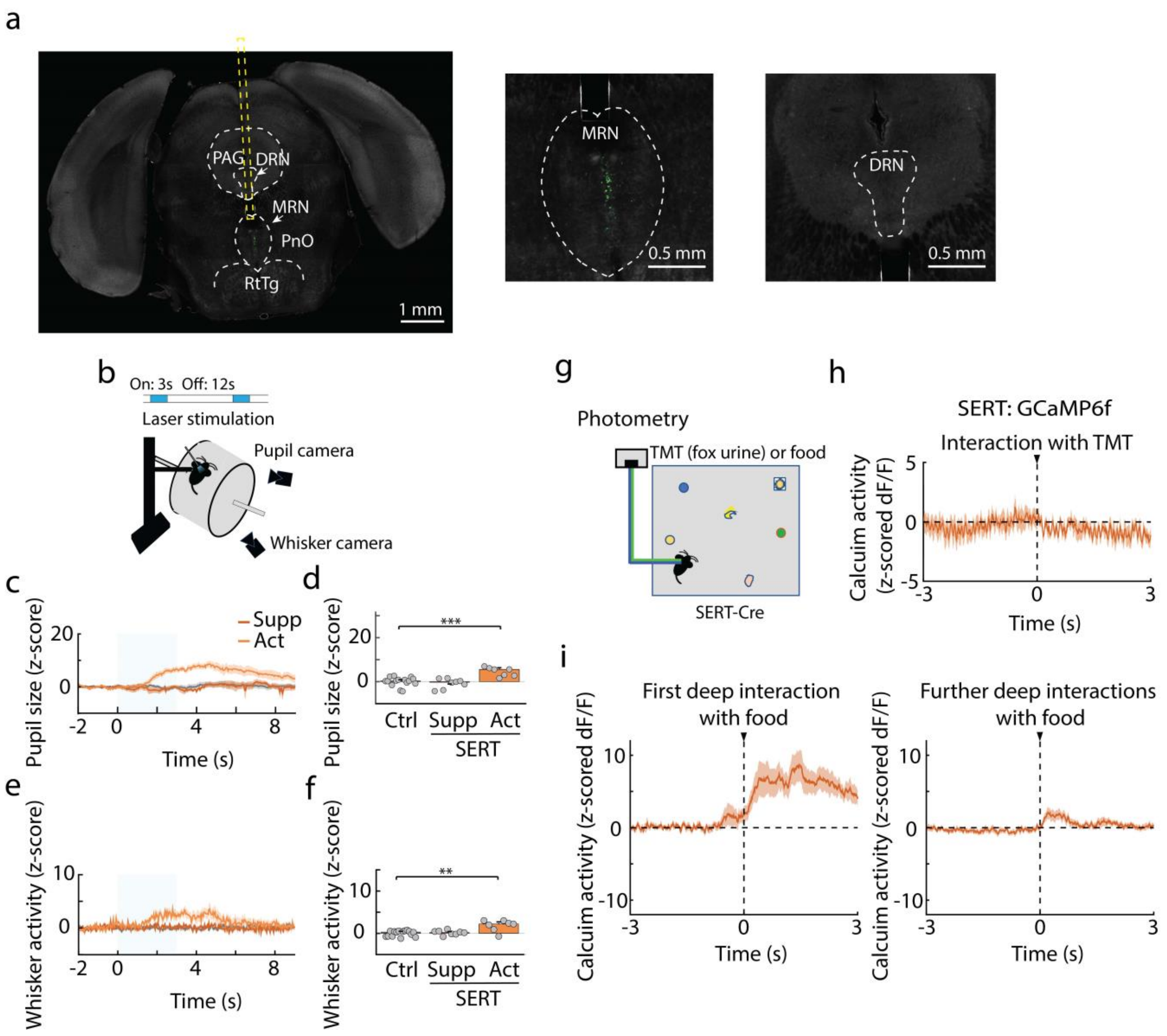


**Supplementary Figure 6. SERT+ MRN neurons do not respond to negative salient stimuli**

**a,** Example image of virus expression (stGtACR2-fusionRed) in SERT+ neurons in the median raphe nucleus (MRN) with optic fiber position (top), with zoomed in sections showing labeled neurons in MRN (bottom left) but not in the dorsal raphe nucleus (DRN, bottom right). **b,** Schematic of the experimental design to examine the effects of optogenetic manipulation of SERT+ MRN neurons on arousal level, measure using pupil size and whisker activity. **c,** Z-scored pupil size over time (mean ± s.e.m.) in control mice (grey) and mice with activation or suppression of the SERT+ MRN neurons, averaged over trials and aligned to onset of optogenetic manipulation. Light blue bar indicates the laser stimulation period. **d,** Median z-score of pupil size during laser stimulation from traces in **c**. N = 15, 8 and 7 mice; $P > 0.9999$ and $P = 0.0004$ for comparing control mice to activation or suppression of SERT+ neurons, respectively; two-sided t-test with Bonferroni multi-comparison correction. **e,f,** same as **c,d** but for z-scored whisker activity (summation of absolute frame-by-frame differences in pixel luminance). N = 15, 8 and 7 mice, respectively; $P > 0.9999$ and $P = 0.0010$ for comparing control mice to activation or suppression of SERT+ neurons, respectively; two-sided t-test with Bonferroni multi-comparison correction. **g,** Schematic of the fiber photometry recording during interactions with food or a TMT-coated object. **h,** Average z-scored calcium trace (mean ± s.e.m., n = 4 mice) of SERT+ MRN neurons aligned to the onset of interactions with an aversive, TMT-covered object. **i,** Average z-scored calcium trace of SERT+ MRN neurons aligned to the first deep interaction and further deep interactions with a food pellet. **: p-value < 0.01, ***: p-value < 0.001.

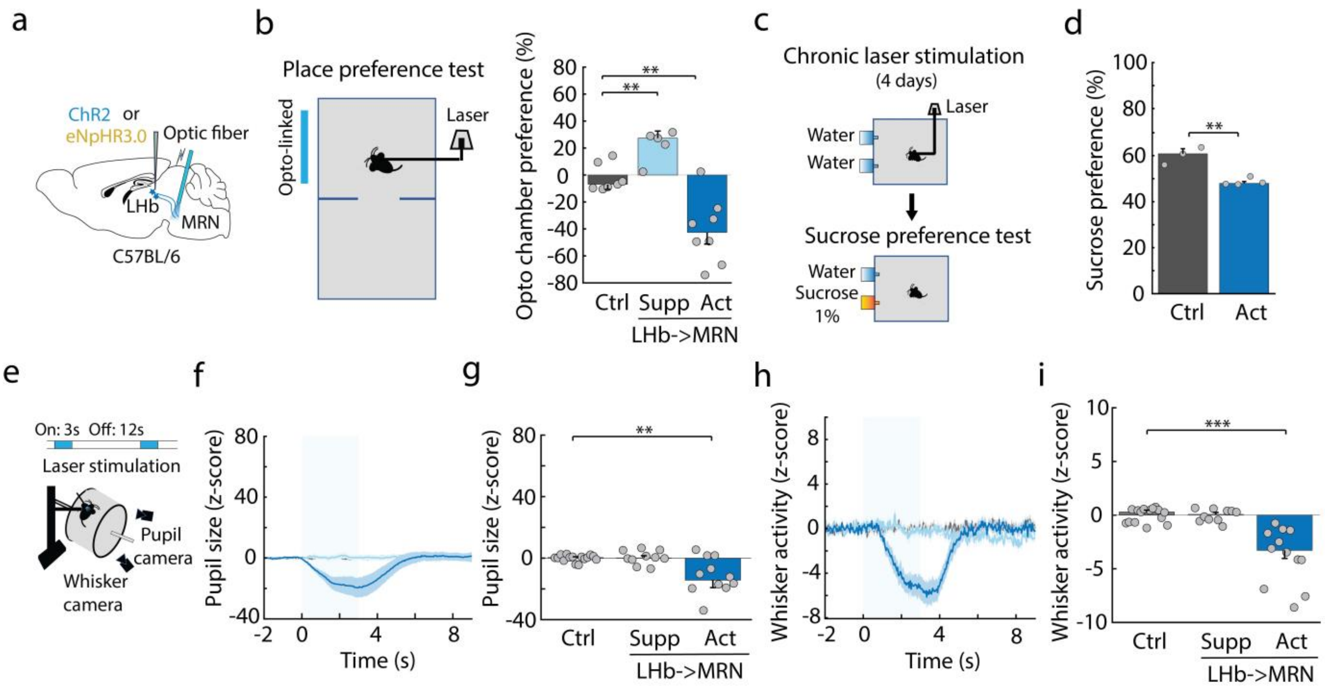


**Supplementary Figure 7. Impact of manipulation of LHb input to MRN on valence and arousal**

**a,** Schematic of experimental design for optogenetic suppression or activation of LHb input to MRN. **b,** Left: Schematic of the real-time place preference test. Right: preference for the opto-linked chamber (100 × (duration of time spent in the opto-linked chamber - duration of time spent in the non-stimulation chamber) / total time) in control mice and mice with activation or suppression of LHb input to MRN (N = 7, 5 and 8 mice, respectively; P = 0.0053 and P = 0.0034 for comparing control mice to activation or suppression of LHb input to MRN, respectively; two-sided t-test with Bonferroni multi-comparison correction). **c,** Schematic of the experimental design: chronic laser activation (4 days, 24 hours per day) followed by the sucrose preference test without laser stimulation on the 5th day. **d,** Sucrose preference in control mice and mice after repeated optogenetic activation of LHb input to MRN (N = 3 and 4 mice, respectively, P = 0.0027, two-sided t-test). Bars depict median, error bars are bootstrapped standard error and circles indicate individual experiments. **e,** Experimental design to examine the effects optogenetic manipulation of LHb input to MRN on arousal level, measure using pupil size and whisker activity. **f,** Z-scored pupil size over time (mean ± s.e.m.) in control mice and mice with suppression or activation of LHb input to MRN, averaged over laser stimulation trials and aligned to laser onset. The light blue bar indicates the laser stimulation period. **g,** Median z-scored pupil size during the laser stimulation period from traces in **f**. N = 15, 10 and 12 mice; P > 0.9999 and P = 0.0037 for comparing control mice to suppression or activation of LHb input to MRN, respectively; two-sided t-test with Bonferroni multi-comparison correction. **h,i,** same as **f,g** but for z-scored whisker activity (summation of absolute frame-by-frame differences in pixel luminance). N = 15, 10 and 12 mice; P > 0.9999 and P = $2.4\times10^{-5}$ for comparing control mice to suppression or activation of LHb input to MRN, respectively; two-sided t-test with Bonferroni multi-comparison correction. **: p-value < 0.01, ***: p-value < 0.001.

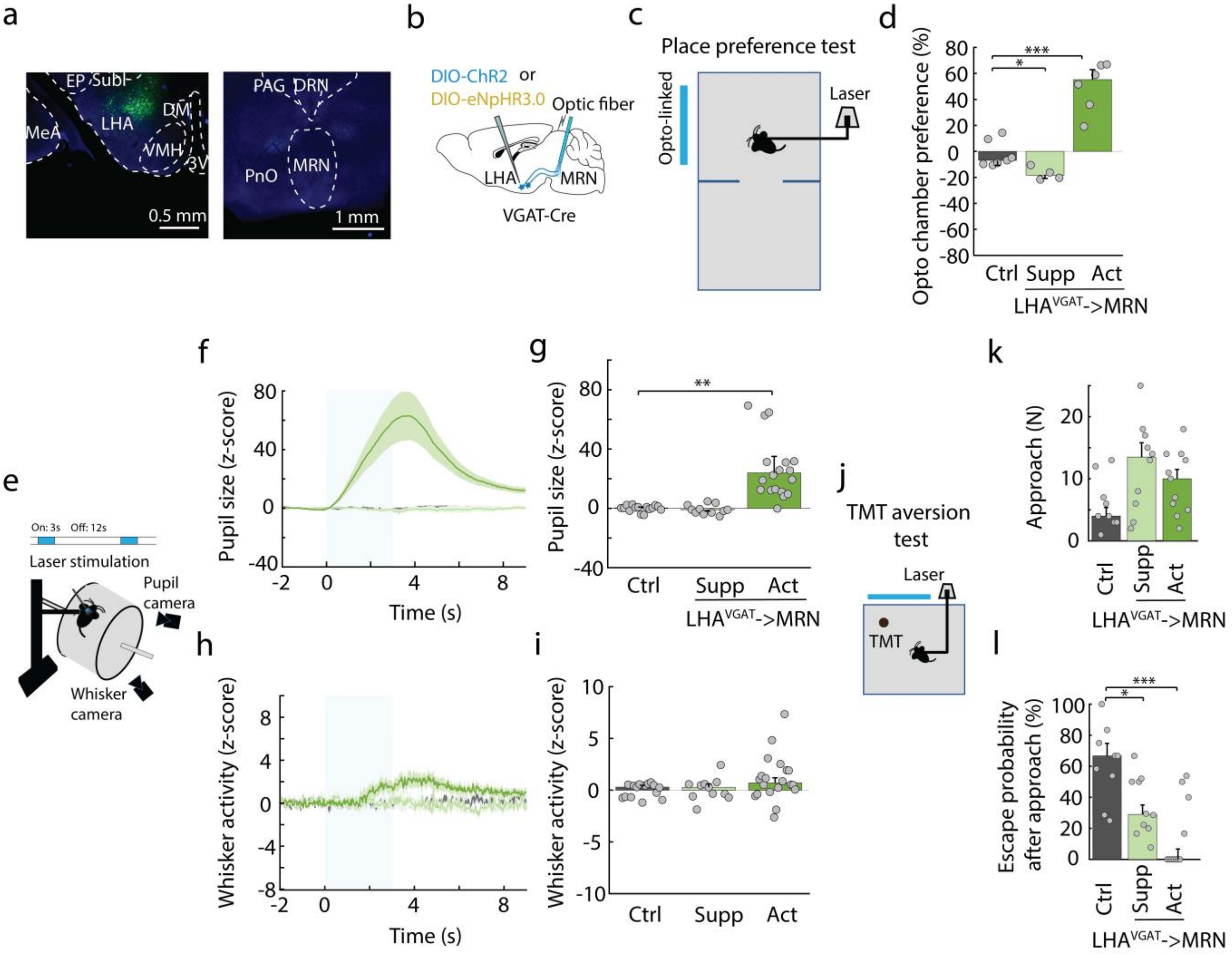


**Supplementary Figure 8. Impact of manipulation of LHA VGAT+ input to MRN on valence and arousal**

**a,** Example image of virus expression in VGluT2+ neurons in LHA (in green; left) of VgluT2-Cre mice, showing absence of axon terminals in MRN (right). 3V: 3rd ventricle, DM: dorsomedial hypothalamus, EP: entropeduncular nucleus, LHA: lateral hypothalamic area, MeA: medial amygdala, Subl: subincertal nucleus, VMH: ventromedial hypothalamus. DRN: dorsal raphe nucleus, MRN: median raphe nucleus, PAG: periaqueductal gray, PnO: pontine reticular formation. **b,** Schematic of experimental design for optogenetic suppression or activation of LHA VGAT+ input to MRN. **c,** Schematic of the real-time place preference test. **d,** Preference for the opto-linked chamber (100 × (duration of time spent in the opto-linked chamber - duration of time spent in the non-stimulation chamber) / total time) in control mice and mice with activation or suppression of LHA VGAT+ input to MRN (N = 7, 4 and 6 mice, respectively; P = 0.04865 and P = 0.0001 for comparing control mice to activation or suppression of LHA VGAT+ input to MRN, respectively; two-sided t-test with Bonferroni multi-comparison correction). **e,** Experimental design to examine the effects of optogenetic manipulation of LHA VGAT+ MRN input on arousal level, measured using pupil size and whisker activity. **f,** Z-scored pupil size over time (mean ± s.e.m.) in control mice and mice with suppression or activation of LHA VGAT+ input to MRN, averaged over laser stimulation trials and aligned to laser onset. The light blue bar indicates the laser stimulation period. **g,** Median z-scored pupil size during the laser stimulation period from traces in **f**. N = 15, 11 and 20 mice; P > 0.9999 and P = 0.0044, for comparing control mice to suppression or activation of LHA input, respectively; two-sided t-test with Bonferroni multi-comparison correction. **h,i,** same as **f,g** but for z-scored whisker activity (summation of absolute frame-by-frame differences in pixel luminance). N = 15, 11 and 20 mice; P > 0.9999 and P = 0.1352 for comparing control mice to suppression or activation of LHA input, respectively; two-sided t-test with Bonferroni multi-comparison correction. **j,** Schematic of the TMT aversion test. **k,** Number of approaches of the TMT-covered object in control mice and mice with suppression or activation of LHA VGAT+ input to MRN. N = 9, 10 and 11 mice, respectively; P = 0.0760 and P = 0.2107 for comparing control mice to suppression or activation of LHA input, respectively; two-sided t-test with Bonferroni multi-comparison correction. **l,** Escape probability after approaching the TMT-covered object for mice shown in **k**. P = 0.0258 and P = 0.0005 for comparing control mice to suppression or activation of LHA input, respectively; two-sided t-test with Bonferroni multi-comparison correction. *: p-value < 0.05, **: p-value < 0.01, ***: p-value < 0.001. In panels **d**, **g**, **i**, **k** and **l**, bars depict median, error bars are bootstrapped standard error and circles indicate individual experiments.

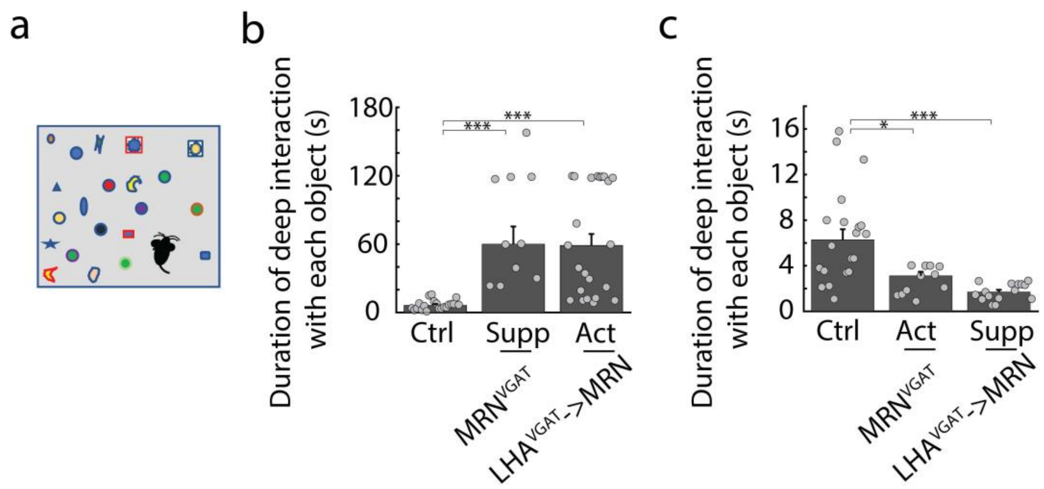


**Supplementary Figure 9. Manipulation of LHA VGAT+ input to the MRN and of VGAT+ MRN bidirectionally regulates the duration of deep interactions**

**a,** Schematic of the MNOI test. **b,** Duration of deep interactions with each object during the MNOI test in control mice (ctrl), mice with suppression of VGAT+ MRN neurons (supp. vgat) and mice with activation of LHA VGAT+ input to the MRN (act. lha). Ctrl vs. supp. vgat: $P = 1.6\times10^{-6}$, ctrl vs. act. lha: $P = 7.6\times10^{-6}$, two-sided t-test with Bonferroni multi-comparison correction. N = 20, 10 and 23 experiments from 10, 5 and 9 mice in ctrl, supp. vgat and act. lha groups. **c,** Duration of deep interactions with each object in the MNOI test in control mice (ctrl), mice with activation of VGAT+ MRN neurons (act. VGAT) and mice with suppression of LHA VGAT+ input to the MRN (supp. LHA), Ctrl vs. act. VGAT: P = 0.0102, ctrl vs. supp. LHA: P = 0.0001, two-sided t-test with Bonferroni multi-comparison correction. N = 20, 11 and 15 experiments from 10, 6 and 8 mice in ctrl, act. VGAT and supp. LHA groups. *: p-value < 0.05, p-value < 0.001. Bars depict median, error bars are bootstrapped standard error and circles indicate individual experiments.

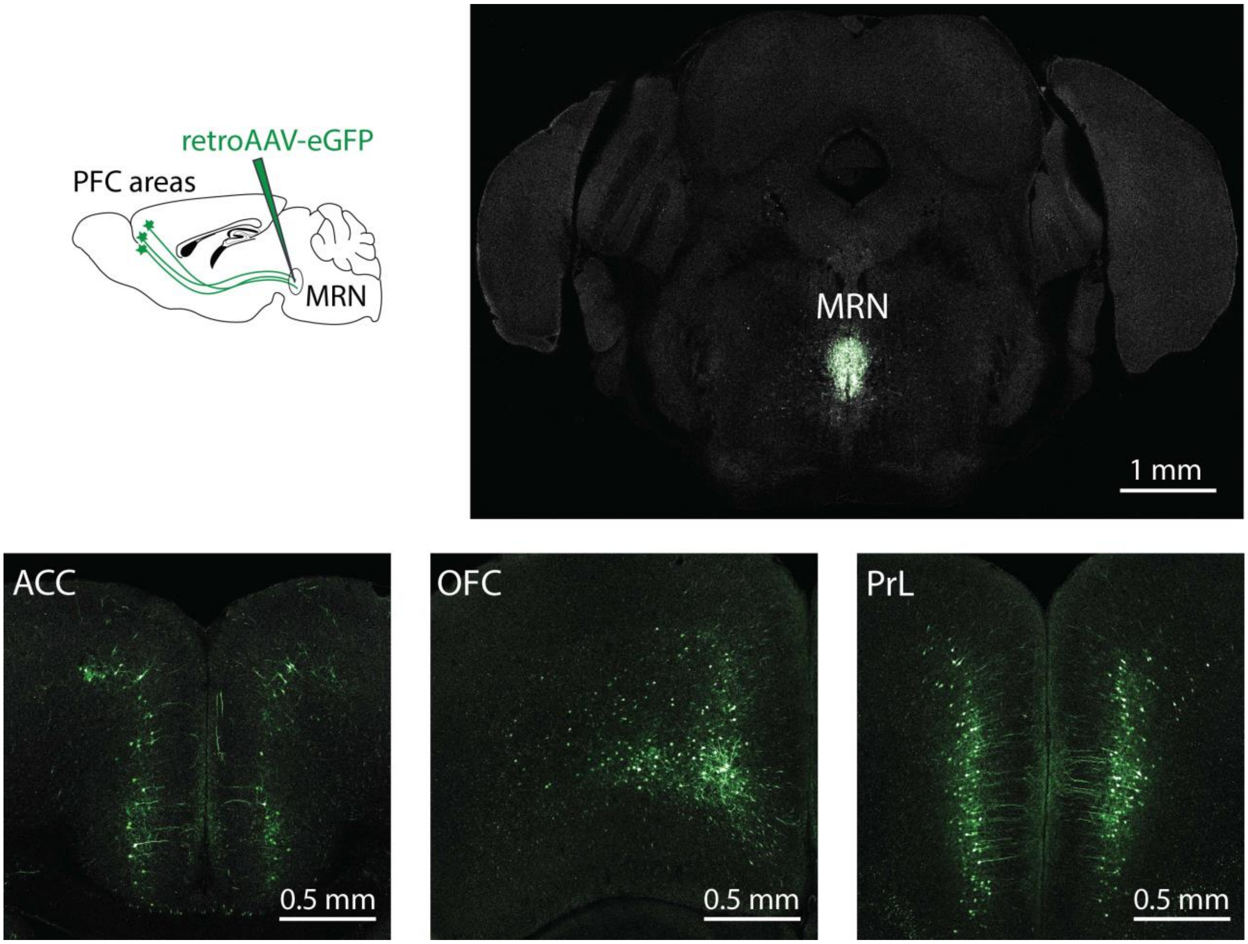


**Supplementary Figure 10. PFC inputs to MRN**

Schematic of retrograde tracing from MRN using retrograde AAV, an example image of virus expression in the MRN (top) and MRN-projecting prefrontal cortical neurons (bottom) in ACC, OFC, and PrL. ACC: anterior cingulate cortex, OFC: orbitofrontal cortex, PrL: prelimbic cortex.
